# Using high-dimensional immunophenotyping for mapping immune cell composition in rapid cryopreserved liquid biopsies

**DOI:** 10.1101/2025.01.13.632307

**Authors:** JG Pedersen, JCL Jordt, E Thorborg, LL Dietz, A Winther-Larsen, KR Gammelgaard, MR Jakobsen

## Abstract

Liquid biopsies, such as a blood sample, are non-invasive and simple ways to monitor biomarkers in clinical settings. For cancer immunotherapy settings a major clinical goal is to identify host immunological biomarkers that can be used to monitor progression of disease, treatment efficacy, or even adverse effects. Here, the profiling and validation of blood circulating immune cells may allow continuously survey on how the immune system alters during disease progress and treatment. However, such practice will require proper sampling and development of a robust high-dimensional phenotypic analysis.

Here, we designed a 37-marker spectral flow cytometry assay with the purpose of conducting high-dimensional immunophenotyping as monitoring the immune status in patients. The assay was developed to profile all major immune cell compartments, their differentiation as well as activation markers, including numerous checkpoint markers. Importantly, we validated this assay in context of how liquid biopsies should be processed to ensure minimal effect of cryopreservation compared to immunophenotyping of a freshly acquired blood sample. Our work highlights important considerations for which phenotypic alternations can be introduced with different cryopreservation methods. Essentially, we identified a fast and simple processing pipeline where cryopreservation of immune cells showed minimal changes in the immunophenotyping profile. This approach can certainly be implemented in clinical settings and will help diagnostic clinical sites with a strong and validated tool applicable for monitoring individual patients’ immunological status.

## Introduction

The applicability of liquid biopsies in cancer settings for early screening and monitoring of disease progression is slowly becoming a general clinical practice. This is partly due to the non-invasive, simple, and low time-consuming manner of taking a blood sample compared to more traditional tumor tissue biopsies. Liquid biopsies have great potential as they contain numerous relevant biomarkers which can be used to define a specific clinical or therapeutic status of a patient. Since liquid biopsies are less invasive, their use allow clinicians to continuously survey the patient during disease process and treatment, and closely evaluate and adjust treatment regimens to achieve a well-fitted personalized precision treatment schedule [1, 2].

Processing of a liquid blood biopsy is often accompanied by analyzing components in serum or plasma. Both matrixes are applicable to assess secreted cytokines, circulating tumor cells, circulating tumor DNA and other biomarkers. However, a major component of a liquid blood biopsy is the immune cell compartment, which provides detailed information about a patient’s homeostatic condition and disease development. These cells are often neglected in clinical settings due to a complex and cumbersome handling process. Nonetheless, evaluating the phenotypic profile of blood immune cells, their differentiation and activation status holds many possibilities for providing clinicians with information about the immune status of patients prior to and during therapy.

The immune cell composition and phenotypic changes in liquid blood biopsies have formerly been assessed using flow cytometry with various sizes of antibody panels including cell surface markers and intracellular markers. Overall, these assays have been constrained by the number of markers per panel due to challenges with the spectral overlay of individual fluorophores and the technical limitations of conventional flow cytometer. Thus, assessment of large phenotypic panels (exceeding 15-20 markers) has not been feasible. However, recent technological advances such as mass cytometry by time-of-flight (CYTOF) and spectral flow cytometry (SFC) now enable scientists to achieve higher dimensional analysis of immune cells within a liquid biopsy [3, 4]. CYTOF uses antibodies tagged with heavy metal isotopes and thus has essentially no issues with overlapping signals from each marker. However, CYTOF will not allow studying cell size and granularity, and it is a slow processing method taking hours to run a single sample. In contrast, SFC offers a rapid and easy transition from conventional flow cytometry systems. High-end SFC equipment allow panel designs of 40-50 markers with minimal interference between markers, if a proper panel design is applied. This can be achieved as the spectrum of each marker fluorochrome is assessed simultaneously, followed by a mathematical deconvolution algorithm that enables precise separation of each marker, akin to a unique fingerprint for each marker in a panel. Thus, though CYTOF may exceed the total number of markers in a panel, SFC is superior when it comes to sample speed, cell size/granulation and in some cases higher sensitivity for low-abundant proteins.

Various studies have described flow panels designed for evaluating immune cell composition in liquid biopsies from cancer patients. For example, in non-small cell lung cancer the immune cell composition was compared between tumor, lymph node and blood using 6 different antibody panels [5]. In another study, immune cell composition in baseline and follow-up blood samples 3 months after immunotherapy was assessed using a 13-color flow panel [6]. In other cases, more selective immunophenotyping panels have been used to assess explicit immune cell composition such as exhaustion markers on T cells [7, 8]. In melanoma, the possibility of using immunophenotyping of blood samples to correlate efficacy of checkpoint inhibitors and progression/non-progression has also been studied [9]. In all, these studies showcase that monitoring immune cell composition in diagnostic settings can be both time-consuming and with the need of multiple parallel assays and a low-dimensional resolution, which are factors that can be resolved using SFC. Another conceivable obstacle for using high-dimensional immune cell analysis in diagnostic laboratories is the need for preparing and storing samples on an ongoing basis that may allow for batch-based processing and post analysis to a lower time and consumable costs. Such processing pipeline requires specific and consistent handling of liquid biopsies and cryopreservation of cells to minimize the effects on marker expression and viability of cells. Inconsistency in sampling methods due to time-consuming procedures possesses the risks of introducing bias and discrepancy in the obtained phenotypes of the characterization of the immune cells.

Here, we present the development of high-dimensional SFC immune cell panel containing 37 markers designed to cover the majority of circulating immune cells and more specifically, to analyse the level of differentiation and exhaustion phenotype within the blood circulating immune cell compartment. Second, to increase the potential of implementing this assay in clinical settings we further explored and validated different liquid biopsy processing and cryopreservation methods to assess which performed best based on processing time, easiness, cell type frequency, and phenotype character consistency, when such samples were compared to analysis of a fresh blood sample. We find the latter analysis highly relevant as it can determine to which extent cryopreservation alters the characteristics of the immune cell composition.

## Results

### Design of the Immunophenotyping Panel for Spectral Flow Cytometry

The SFC panel was designed to comprehensively capture all major blood-circulating immune cell populations and their differentiation status, determined to be relevant for cancer settings (Figure 1). In the context of studying how processing and cryopreservation affected the cell product, we had a particular interest in also assessing a rather broad range of classical markers used to subdivide lymphocytes and myeloid cell populations (Table 1). We initially used the pan-hematopoietic marker CD45 and granulocyte marker CD66b to separate immune cells from non-immune cells (Figure 1A and Figure S1). We then included lineage markers to gate cells into myeloid (CD14^+^ and CD11c^+^) and lymphoid (CD123^+^, CD3^+^, CD56^+^, CD19^+^) populations. Within the myeloid population we used additional markers to segregate classical (CD14^+^CD16^-^), intermediate (CD14^+^CD16^+^) and non-classical (CD14^-^CD16^+^) monocytes; as well as dendritic cells (DCs) (CD11c^+^HLA-DR^+^). For the lymphoid population, we gated the T cells (CD4 or CD8) into both naïve (CD27^+^CD45RA^+^CCR7^+^), central memory (CD27^+^CD45RA^-^CCR7^+^), terminal effector (CD27^-^CD45RA^+^CCR7^-^), effector memory (CD27^-^CD45RA^-^CCR7^-^) and transitional memory (CD27^+^CD45RA^-^CCR7^-^) cells. Th1 and Treg cells were gated from the CD4+ cluster using Tbet and FoxP3 and CD25, respectively. The NK cells (CD56^+^CD3^-^) were gated based on early (CD56^high^CD16^-^), mature (CD56^high^CD16^high^) and terminal (CD56^dim^CD16^high^) differentiation. For B cells (CD19^+^), we only included markers that allowed gating into naïve (CD27^-^) and memory (CD27^+^) cells. Finally, using CD123 and HLA-DR we could also gate on plasmacytoid DCs (pDCs).

**Figure 1.**
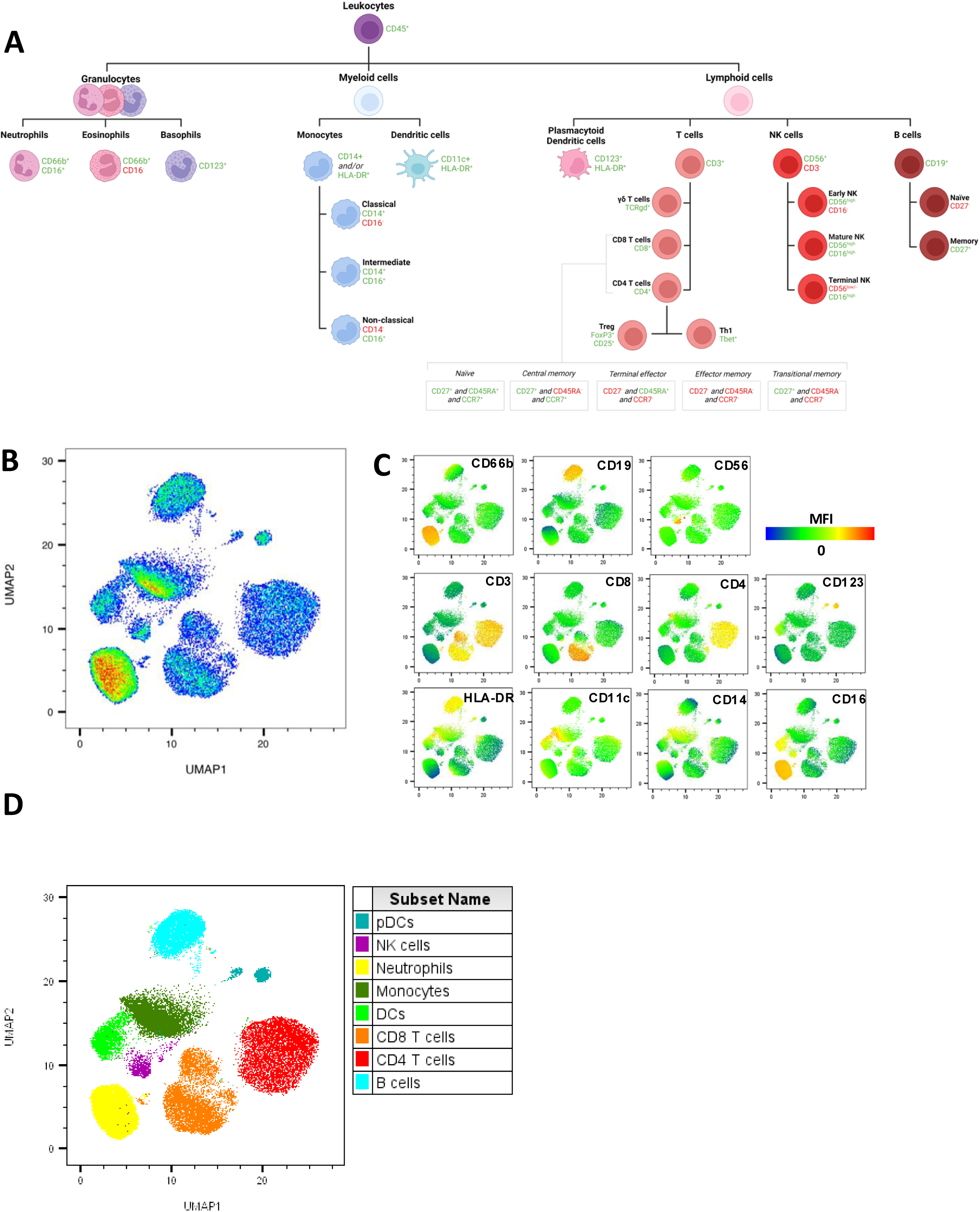
Development of the 37-marker spectral flow cytometry panel. **A)** Illustration of all immune cell types that can be identified within the panel of markers used in the spectral flow analysis. **B)** A representative UMAP plot of a fresh whole blood sample analysed with the 37-markers panel. The UMAP plot was based on pre-gated live CD45^+^ cells and generated using lineage markers (CD66b, CD19, CD56, CD3, CD8, CD4, CD123, HLA-DR, CD11c, CD14, CD16) as input parameters. **C-D)** Lineage marker expression overlay on individual UMAP plots and **D)** UMAP overlaid with clusters generated by the FlowSOM algorithm using lineage markers as input parameters.

**Table 1:**

| Lineage | Differentiation | Activation Lymphoid | Activation Myeloid | Others |
| --- | --- | --- | --- | --- |
| CD45 | CD8 | CD28 | CD86 | Live/dead |
| CD66b | CD4 | CD38 | CD163 | Ki67 * |
| CD3 | HLA-DR | CD69 | PD-L1 |  |
| CD19 | CD27 | CD73 |  |  |
| CD14 | CCR7 | IFNg * |  |  |
| CD56 | CD45RA | ICOS |  |  |
| CD123 | CD11b | PD-1 |  |  |
| CD11c | CD25 | TIGIT |  |  |
|  | Foxp3 * | LAG3 |  |  |
|  | Tbet * | CTLA4 * |  |  |
|  | CD16 | TIM3 |  |  |
|  | TCRgd |  |  |  |
| 8 | 12 | 12 | 3 | 2 |

The 37-marker panel was defined by lineage markers, classical differentiation markers, as well as activation markers for both lymphoid and myeloid immune cells. \*Markers included in the intracellular staining.

All antibodies included in the SFC panel were first mathematically tested for spillover spreading (Figure S2), and then titrated using frozen peripheral blood mononuclear cells (PBMCs) to determine optimal staining concentrations (Figure S3 and S4). Here, we used both single stain and subsequently all-strain and fluorescence minus one (FMO) controls to ensure the compatibility of all 37-markers within the same staining procedure. This work confirmed that most markers including HLA-DR, CD19 and CD11b behaved appropriately in both single stain and all-stain tests (Figure S5A). In contrast, other markers tested demonstrated poor staining quality in the all-stain samples and needed optimization. Among these, the Tbet AF532 (clone S18004B) antibody only worked in the single stain setting but not in the all-stain setting most likely due to a poor antibody clone and a too dim fluorochrome. However, this was resolved by changing to a different antibody clone and a brighter fluorophore (Figure S5B). A general concern with SFC is the risk of data spreading with expanding numbers of markers in the all-stain panel. We observed this for example with the monocyte marker CD14, but again, this could be resolved by switching both the antibody clone and fluorophore (Figure S5C). Following thorough titrations, optimization and validation, we were able to construct a clear distinguishable clustering of cell types within a blood sample using the Uniform Manifold Approximation and Projection for Dimension Reduction (UMAP) algorithm (Figure 1B). Subsequently, we confirmed, based on marker expression intensity, that the clusters were distinct populations representing all the major lymphocyte populations (pDCs, T, B and NK cells), neutrophils/granulocytes, and the major myeloid populations (monocytes and DCs) (Figure 1C and D).

### Validation of Activation and Differentiation Markers

Importantly, our panel included several phenotyping markers which allow us to increase the high-dimensional resolution of individual immune cell types within a rather complexed mixture of cell populations seen in blood (Table 1). The panel contains immune checkpoint markers such as: PD-1, LAG3, TIM3, CTLA4, and TIGIT; Inducible T cell costimulatory ICOS; co-receptor CD28; and cyclic ADP ribose hydrolase CD38. We included CD69 as another early marker for lymphocyte activation as well as intracellular IFNγ staining as an activation marker for both NK, CD4 and CD8 T cells. The CD73 receptor was included as a broad immune checkpoint marker expressed on both B cells, NK, myeloid and T cells [10], and has been reported to be expressed on CD4 Tregs upon immunosuppressive conditions but shows limited expression on healthy CD4+ Tregs. For myeloid activation, we focused on the major activation markers such as PD-L1, CD163 and CD86. To properly validate each of the antibodies targeting these markers on frozen PBMCs, we had to introduce *in vitro* stimulation of cells to allow upregulation of each marker. For this, we used the condition of Phorbol 12-myristate 13-acetate (PMA) plus ionomycin during a 24-hour culturing assay (Figure 2A). Each of the 16 markers were then titrated and validated for single stain and all-stain in each condition (Figure S4). We subsequently, tested the full panel of activation/differentiation markers in combination on unstimulated and PMA/Ionomycin stimulated PBMCs (Figure 2B-C). We first conducted dimensionality reduction analysis using tSNE (Figure 2B) combined with the FlowSOM clustering algorithm on CD45+CD3+ gated cells (Figure 2B). To visualize the changes in MFI for each marker we calculated a normalized z-score value, which was plotted in a heatmap (Figure 2C). From these data, we confirmed that each relevant marker was expressed in either unstimulated or stimulated lymphocytes and/or monocytes.

**Figure 2.**
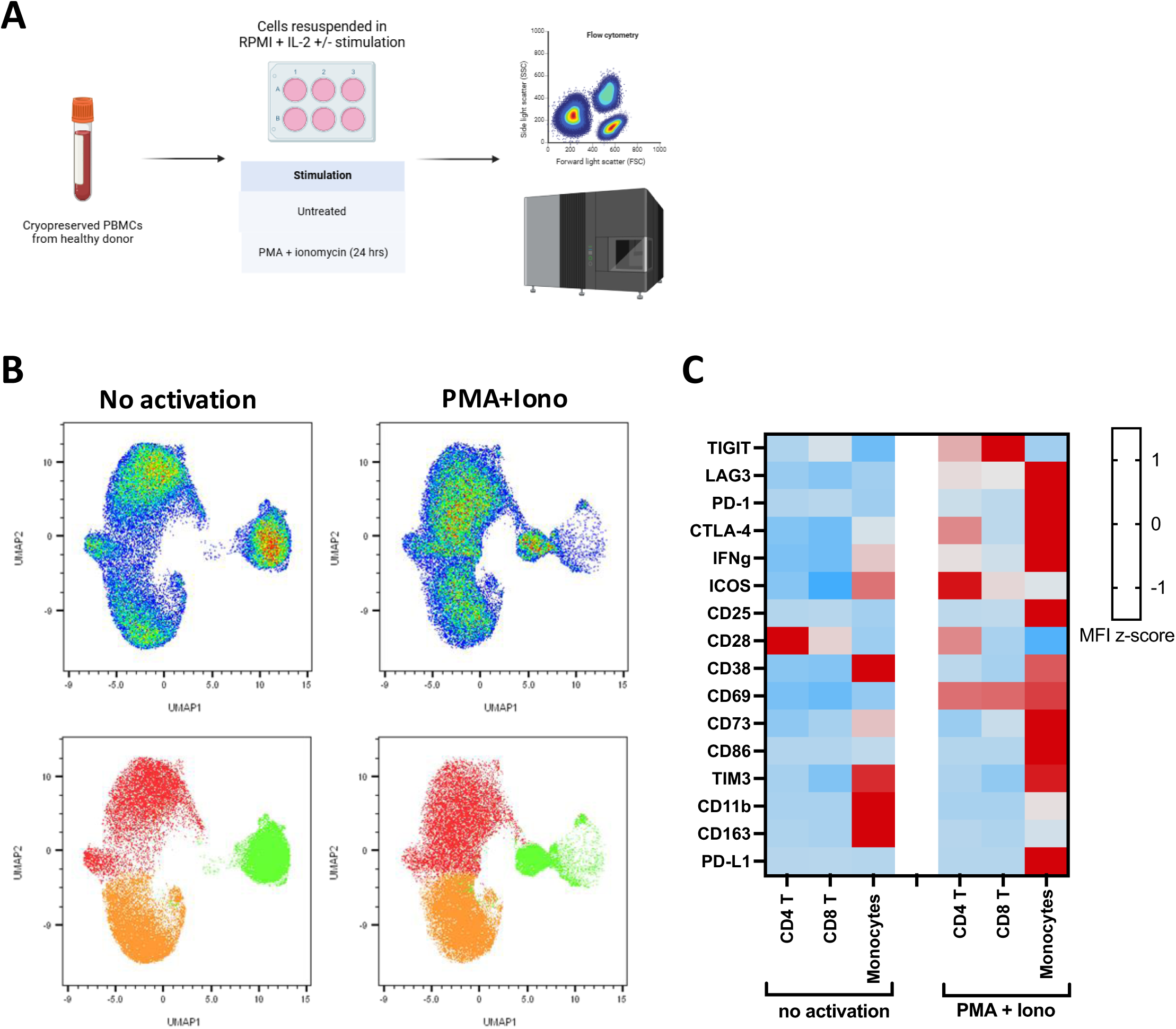
Evaluation of activation and exhaustion markers. **A)** Illustration of the experimental setup for activation of PBMCs. **B)** Representative UMAP plots of a thawed PBMC sample with and without activation analysed with the 37-marker SFC panel. The UMAP plots were generated on a combined lymphoid and myeloid cell cluster (lymphoid: CD45+CD3+, myeloid: CD45+CD3-CD14+) and overlaid with clusters generated by the FlowSOM algorithm using Forward scatter (FSC), Side scatter (SSC), lymphoid (CD3, CD4 and CD8) and myeloid (CD14 and HLA-DR) markers as input parameters. **C)** Heatmaps showing the expression (MFI, median fluorescence intensity) of lymphoid activation and exhaustion markers (TIGIT, LAG3, CD73, CD28, CD69, TIM3, CD25, PD-1, IFNg, CTLA-4, ICOS, CD38) and myeloid activation and exhaustion markers (CD11b, TIM3, CD86, CD163, PD-L1) across activation status in FlowSOM-generated clusters. MFIs were normalized across each marker creating an MFI z-score.

In summary, the 37-marker SFC panel (see Table S1) enables a clear sub-clustering of all the major immune cell populations from a liquid blood biopsy into lineage cell compartments and exploration of their activation and differentiation status, generating a high-dimensional immune cell phenotyping profile.

#### Liquid Biopsy Processing Effects on Immune Cell Distribution

Having validated and optimized the SFC high-dimensional immune cell phenotyping panel (hereafter HD-ICPP), we next wanted to explore whether different handling processes and cryopreservation of liquid blood biopsies would affect the immune cell composition. For this, we designed a 3×3 matrix comparing processing methods and freezing medium conditions (Figure S6). The processing methods used to obtain immune cells were: 1) collection of whole blood plus red blood cell (RBC) lysis, 2) HetaSep isolation, and 3) SepMate-Ficoll density gradient isolation. The HetaSep process is a gentle and less than a 1-hour process where RBCs aggregate in the blood tube allowing separation of nucleated cells from the RBCs. The SepMate-Ficoll process is a more time-consuming and complex procedure for the collection of mononucleated cells from whole blood. Following the three collection methods, samples were split into three different cryopreservation solvents. First, we tested the compound BloodStor 55-5 (BS55-5) which has been designed for stem cell cryopreservation by an animal component-free pre-formulated with 55% (w/v) DMSO and 5% (w/v) Dextran-40. This cryopreservation reagent can be mixed directly with whole blood, and then frozen – demonstrating the fastest, easiest and most untangled method to handle a liquid biopsy. Additionally, we included two other freezing mediums that are often used in conjunction to ficoll-gradient processed blood, namely standard freezing medium (SFM) formulated with 10% DSMO and 90% heat-inactivated Fetal calf serum; and CryoStor CS10, an animal component-free solvent pre-formulated with 10% DMSO supported to be valued for cryopreservation of sensitive cell types. It is important to notice that samples using whole blood cryopreservation did include RBCs and required a lysis step of the BS55-5-cryopreserved cells after thawing and prior to running the HD-ICPP. For the two other freezing conditions, we had to introduce the RBC lysis step prior to cryopreservation (Figure S6). For all cryopreservation assays, we used the same standard Mister Frosty container to control freeze-rate. After 24 hours at −80°C, samples were moved to a −150°C freezer and stored for at least 7 days before thawing and processed for flow staining.

First, we addressed whether blood sample processing affected immune cell composition. This was done by comparing the most untouched sample – being the fresh whole blood – with each of the two other methods used for PBMC purification from whole blood. We conducted HD-ICPP on blood cells obtained from healthy donors (n=6), followed by unsupervised UMAP and FlowSOM analysis performed by the FlowJo^TM^ software (version 10) for clustering and visualization. A representative unsupervised UMAP clustering from one donor on each processing condition is provided (Figure 3A top panel) with FlowSOM clusters overlayed (Figure 3A bottom panel). Overall, each processing method gave various immune cell composition profiles, which was also donor dependent (Figure 3B). The distribution of cell populations in the fresh samples were divided in 34.65% CD4+ T cells [29.87% (HetaSep); 35.17% (Ficoll)], 28.43% monocytes [34.38% (HetaSep); 25.80% (Ficoll)]; 15.88% CD8+ T cells [13.50% (HetaSep); 16.62% (Ficoll)]; 7.32% NK cells [8.82% (HetaSep); 8.90% (Ficoll)]; 5.88% B cells [5.89% (HetaSep); 6.8% (Ficoll)], 1.67% DCs [1.83% (HetaSep); 1.7% (Ficoll)], 0.53% pDCs [0.61% (HS); 0.64% (Ficoll)]. Notably, we did observe statistically significant difference in the frequency of CD4, CD8 and monocytes when fresh blood or Ficoll was compared to HetaSep, whereas fresh blood and Ficoll gave similar numbers. This was however not reflected in correlation analysis between Fresh whole blood and HetaSep, where there was strong alignment between each cell type on the individual donor level (Figure S7). When we conducted a subcellular analysis within the T cell subpopulations, we did not find any major differences between the three methods in terms of the frequency of effector, terminal, central or transitional memory cells, nor naïve T cells (Figure 3C-D). Similar frequencies were observed for B cells, monocytes and NK cells (Figure 3E-G). As expected, the only major difference between the processing methods was that the granulocytes disappeared when using SepMate-Ficoll (Figure 3A, yellow population).

**Figure 3.**
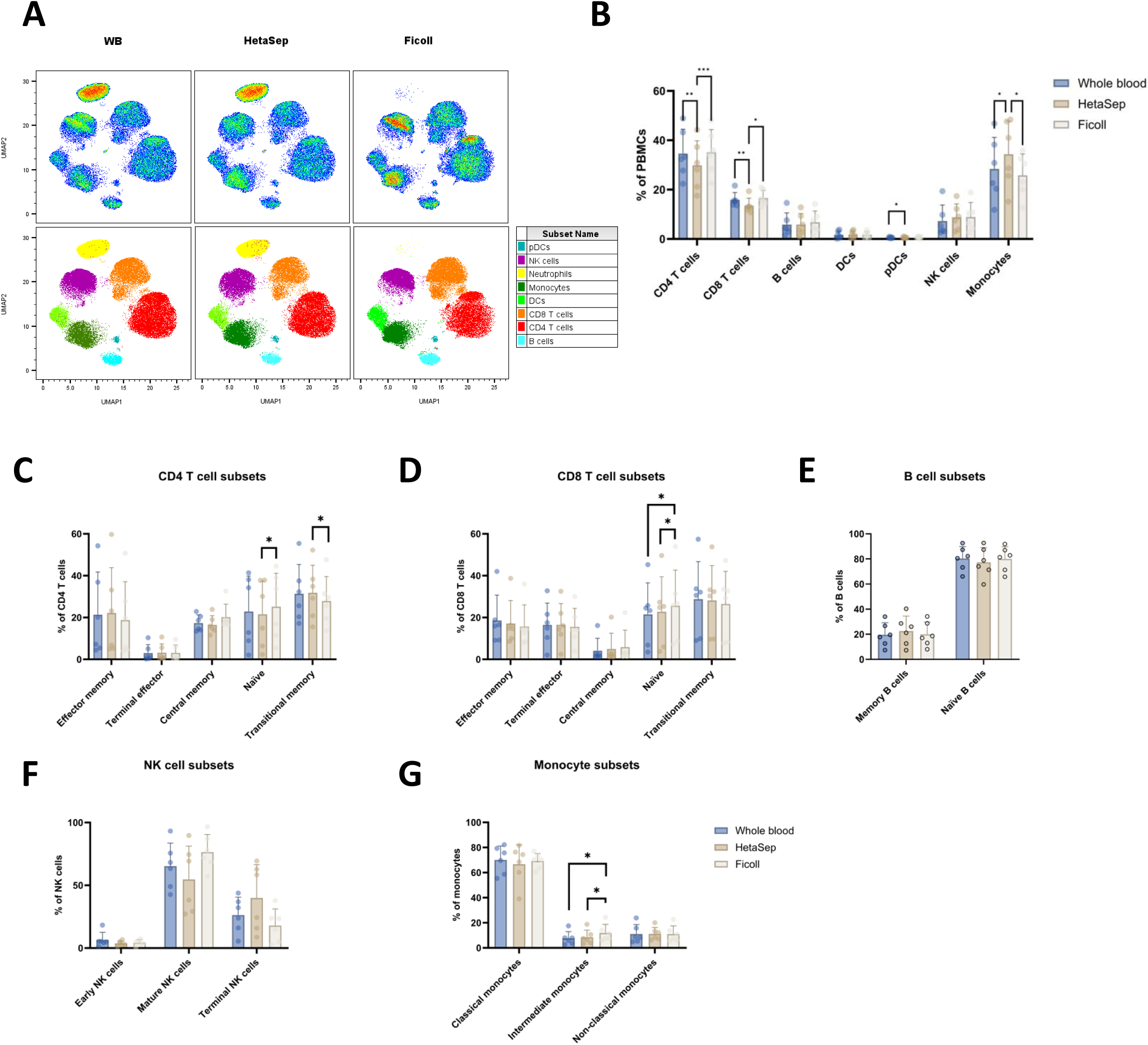
Isolation method effect on PBMC composition in fresh blood samples. **A)** UMAP plots showing the composition of immune cells in freshly collected blood with or without the use of HetaSep and Ficoll processing. The UMAP dimension reduction was computed on live CD45+ cells with lineage markers (CD3, CD4, CD8, CD11c, CD14, CD16, CD19, CD56, CD66b, CD123, and HLA-DR) as input parameters. Overlay plots show immune cell types identified by FlowSOM using the same lineage markers as input parameters. **B)** The Immune cell type frequencies based on manual gating, where frequencies were calculated as percentage of total PBMCs across 6 donors. Statistical significance was tested using a RM one-way ANOVA comparing each isolation method (whole blood (blue), HetaSep (brown), Ficoll (grey)); \**P* < 0.05, *\*\*P* < 0.01, *\*\*\*P* < 0.001. Data presented as dot plots with bars indicate mean values +/- SD as whiskers. **C-G)** Accumulated data across all donors for specific immune cell subset. Frequencies were calculated as percentage of the respective parent immune cell type. The statistical significance was tested using RM One-way ANOVA with Bonferroni’s multiple comparison test comparing each isolation method; \**P* < 0.05, *\*\*P* < 0.01, *\*\*\*P* < 0.001. Data presented as dot plots with bars indicate mean values +/- SD as whiskers.

In conclusion, these data support that no major variations in immune cell composition, including different immune cell subsets, are observed in the three different processing methods for fresh blood samples.

#### Liquid Biopsy Cryopreservation Effects on Immune Cell Distribution

In continuation of our evaluation, we next conducted the HD-ICPP analysis on liquid biopsy samples that had been processed by each of the three methods and subsequently cryopreserved for a minimum of 7 days. From the UMAP analysis, it was apparent that granulocytes did not survive any of the freezing conditions (Figure 4A-B), but besides this, no other obvious differences were observed between fresh and cryopreserved samples, nor between the different cryopreservation methods. Next, to explore the effects of cryopreservation, we decided to use the immunophenotype profile from the whole blood as a reference to each of the processed samples across our donors. Importantly, each of the cryopreservation conditions did not display any significant differences in the immune cell composition across donors as well as across the processing methods (Figure 4C, S9A, S10A). However, when we conducted correlation analysis within each group, we observed a general pattern that the frequency of monocytes differed the most between fresh and each of the processed and cryopreserved methods (Figure S8, S9B-C and S10B-C). Across the data, we observed that CS10 cryopreservation gave the lowest variation within the different cell populations. A major difference between BS55-5 and CS10 cryopreservation that could affect the results, was the use of RBC lysis prior to freezing for CS10, whereas BS55-5 was mixed directly with whole blood and then cryopreserved.

**Figure 4:**
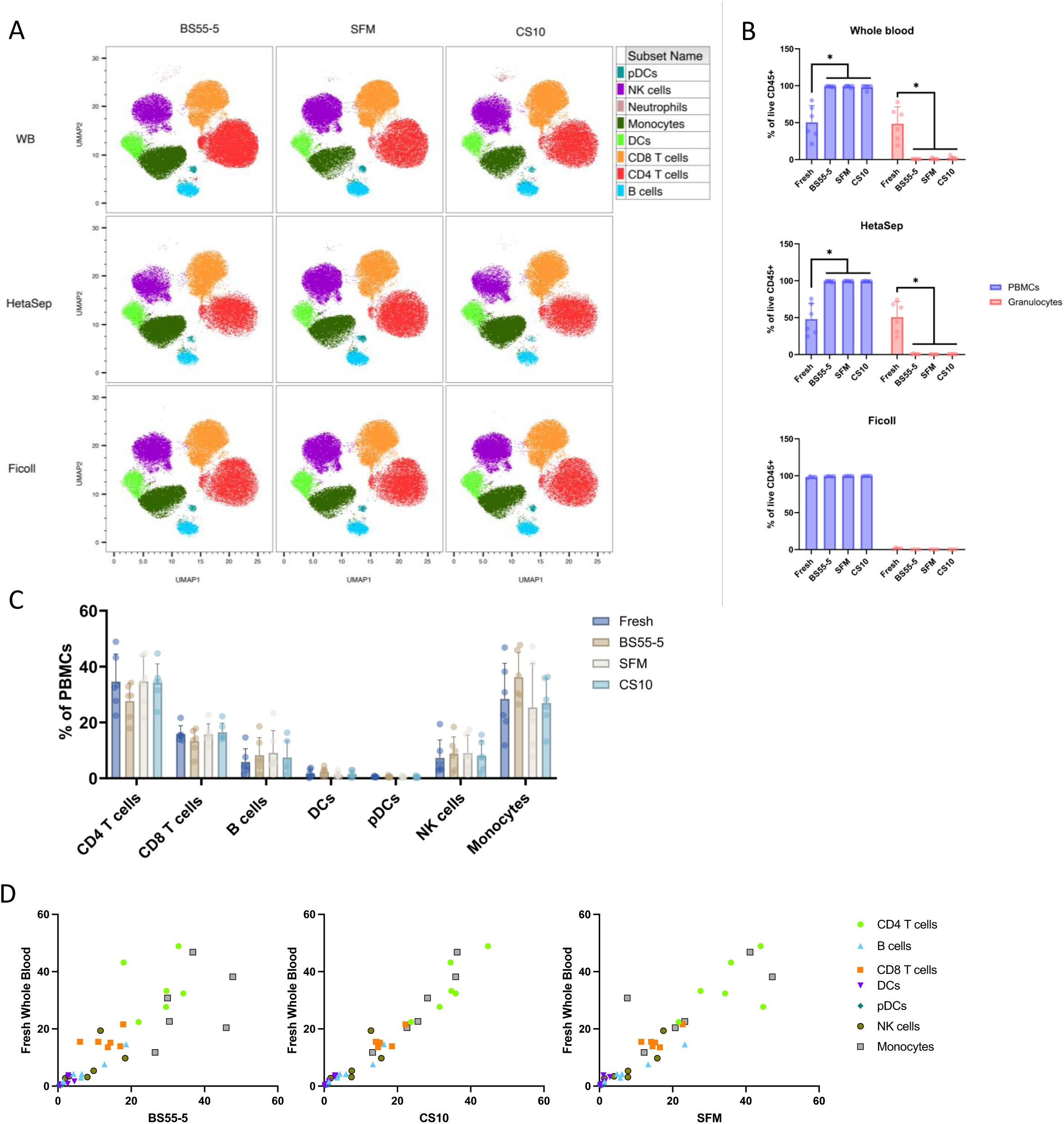
Granulocytes lost during cryopreservation steps. **A)** UMAP plots illustrating the composition of immune cells in the three different cryopreserved (BS55-5, SFM, and CS10) samples from either whole blood, HetaSep or Ficoll processed blood. The UMAP dimension reduction was computed on live CD45+ cells with lineage markers (CD3, CD4, CD8, CD11c, CD14, CD16, CD19, CD56, CD66b, CD123, and HLA-DR) as input parameters. Data is represented from the same donor as used in figure 3A. **B)** Bar graph showing the frequency of PBMCs and granulocytes (as % of live CD45+) in fresh and cryopreserved (BS55-5, SFM, and CS10) samples. Statistical significance was tested using a RM one-way ANOVA test (n = 6) with Bonferroni’s multiple comparison test. \**P* < 0.05. Bars indicate mean with SD as whiskers and individual donors as dots. **C)** Bar graph showing the frequency of major immune cell subsets (as % of live CD45+CD66b-cells) in fresh and cryopreserved (BS55-5, SFM, and CS10) whole blood samples. Statistical significance was tested using a RM one-way ANOVA test (n = 6) with Bonferroni’s multiple comparison test. \**P* < 0.05. Bars indicate mean with SD as whiskers and individual donors as dots. **D)** Correlation analysis between immune cell type frequencies (as % of PBMCs) in fresh whole blood as compared to each of the cryopreservation method (BS55-5, SFM, and CS10). Statistical analysis can be seen in supplementary figure S8.

In conclusion, our results regarding distribution of cell frequency indicate that the most rapid and easy processing method consisting of drawing a whole blood sample, mixing it with CS10 or BS55-5 before cryopreservation performed better than any of the more time-consuming processing methods. Importantly, for all the cryopreservation methods we did observe minor changes in particularly for the monocyte populations besides the anticipated loss of granulocytes.

#### Cryopreservation Effects on Cell Activation Status

It has been suggested that the differentiation and activation status of a particular cell type may impact fragility and sensitivity of the cells to the choice of cryopreservation matrix. To investigate this further, we compared the frequencies of subcellular populations, as determined by differentiation markers, identifiable using the HD-ICPP in both fresh and frozen samples. For the visualization of subcellular CD4 and CD8 populations we decided to conduct a t-distributed stochastic neighbor embedding (tSNE) analysis since this method better preserves inter-cluster relations compared to UMAP. The tSNE and FlowSOM clustering from one donor showed no major differences between the distribution of Terminal, Naïve, Transitional-, Effector- or Central-memory CD4 or CD8 T cells (Figure 5A).

**Figure 5:**
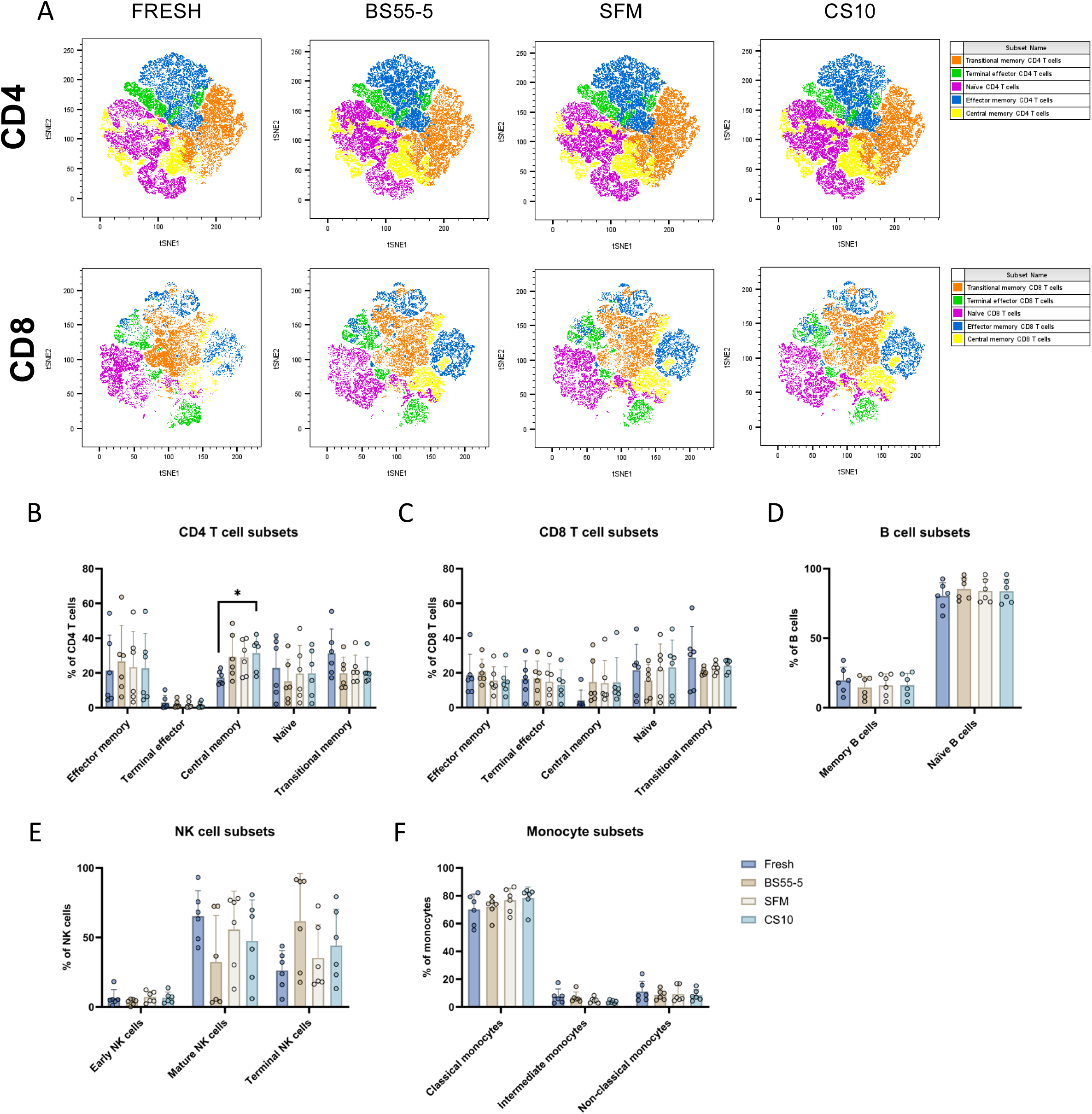
Cryopreservation inflicts only minor changes to the immune cell subset composition. **A)** Dimensionality reduction analysis with tSNE on manually gated CD4 and CD8 T cells from a representative fresh and cryopreserved whole blood sample. The tSNE analysis was computed using CD27, CD45RA, and CCR7 as input parameters for both CD4 and CD8 T cells and clustering of CD4 and CD8 T cell subsets was conducted using the FlowSOM algorithm. **B-F)** Accumulated data from the 6 individual donors, depicturing the changes in frequencies of immune cell subsets **B)** CD4 T cells, **C)** CD8 T cells, **D)** B cells, **E)** NK cells, and **F)** Monocytes. Each cryopreserved sample was compared to their fresh whole blood sample counterpart. The CD4 and CD8 T cells were further divided into effector memory, terminal memory, central memory, naïve, and transitional memory subsets (see also figure 1A), whereas monocytes were divided into Classical, Intermediate, and Non-classical; and NK cells were divided into Early, Mature, and Terminal. Finally, B cells were divided into Memory and Naïve. The statistical significance was tested using a RM one-way ANOVA with Bonferroni’s multiple comparison test including each cryopreserved condition to the reference being the fresh blood sample. \**P* < 0.05, *\*\*P* < 0.01.

Analysis across all donors, showed only minor differences in the composition of CD4 and CD8 T cell subsets between fresh and cryopreserved whole blood samples (Figure 5B-C). We observed changes in the central memory and transitional memory CD4 T cells, with a generally higher frequency of central memory and lower frequency of transitional memory in cryopreserved samples (Figure 5B). No differences between fresh and cryopreserved samples were identified within the various subsets of CD8 T cells (Figure 5C). Next, we conducted a similar analysis on B cell subsets (Figure 5D), NK cell subsets (Figure 5E), and monocyte subsets (Figure 5F), where no significant difference in the frequency was observed. Of note, we did observe a larger variation of each sample points in the NK cell subset suggesting some effects of individual cryopreservation methods used (Figure 5E). To investigate whether cryopreservation introduced stress and other modulatory factors to the cells’ phenotypic characteristics of the cells, we finally evaluated the expression pattern of our 16 pre-defined activation markers using the measured median fluorescent intensity (MFI) level of each marker in the fresh whole blood samples and compared with the different processing and freezing conditions (Figure 6 and Figure S11-13). Overall, freshly collected and stained cells expressed activation markers CD73, CD28, TIM3, TIGIT, CD38, CD11b, CD86, CD163, and PD-1. Both CD4 and CD8 T cells had high expression of CD28 and low expression for PD-1 (Figure 6A; S11A-B); NK cells expressed both CD28, TIGIT, CD38 and CD11b (Figure 6A); B cells had high CD73 expression (Figure 6A); and monocytes a high expression of CD11b, low expression of CD86, CD163, TIM3 and no expression of PD-L1 (Figure 6A; S11C). Next, comparing these expression levels to each of the cell populations from the cryopreserved methods, the most distinct difference was observed based on the CD28 expression. Across all cryopreservation conditions, CD28 expression significantly dropped within all compartments of CD4 and CD8 T cell subsets (Figure 6B-C) and this was independent of the processing methods (Figure S12A-B and S13A-B). For the NK cells, we observed a trend that marker expression patterns were preserved but at the same time a clear drop in both CD38 and TIGIT expression when samples were compared to whole blood (Figure 6D, S12C and S13C). Finally for the monocyte population, we found no significant alternations in marker expression for cryopreserved whole blood (Figure 6E and S11C), but for both HetaSep and Ficoll, a significant drop in TIM3 and CD163 expression was identified (Figure S12D-S13D).

**Figure 6:**
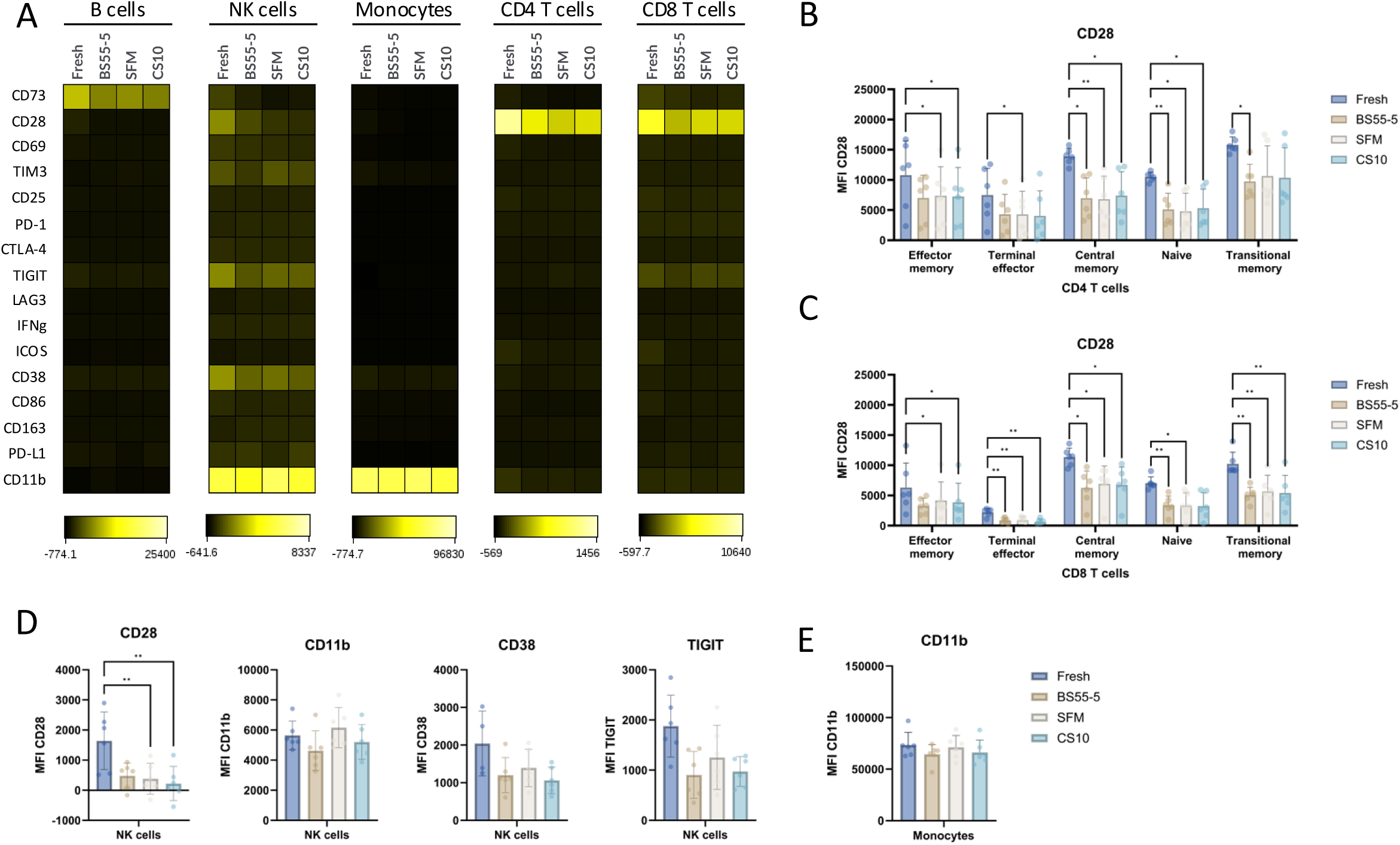
Reduced CD28 expression on T cells and NK cells after cryopreservation. **A)** Heatmap showing the expression levels (MFI, median fluorescence intensity) of each indicated marker across fresh and cryopreserved samples (BS55-5, SFM, and CS10) whole blood samples for B cells, NK cells, monocytes, CD4 T cells, and CD8 T cells, respectively. Each data point represents the median MFI for all 6 donors. **B-C)** Sub-group analysis displaying CD28 expression (MFI) on CD4 T cell subsets **(B)** and CD8 T cell subsets **(C)** across fresh and cryopreserved samples. **D)** Expression of CD28, CD11b, CD38, and TIGT on NK cells from fresh and cryopreserved samples. **E)** CD11b expression on monocytes from fresh and cryopreserved samples. Statistical significance was tested using a RM and one-way ANOVA (B-E) with Bonferroni’s multiple comparison test comparing each cryopreserved condition with fresh sample. \**P* > 0.05; *\*\*P* > 0.01.

In conclusion, our work highlights the importance of monitoring changes for certain activation and differentiation markers, which appear to be impacted by cryopreservation. We showed that this is highly dependent on the processing method and less dependent on the cryopreservation matrix selected. Including parameters such as handling and process time, we find that whole blood sampling, RBC lysis and CS10 cryopreservation was the most suitable method to reflect the landscape of immune cell distribution, differentiation and activation status from a fresh collected blood sample.

## Discussion

In recent years, several studies have highlighted the utility of both CYTOF and SFC for high-dimensional immune cell profiling, often conducted using readily available frozen PBMC samples. These methods hold great promise as monitoring tools for future diagnostics. However, limited attention has been given to the labor-intensive processes required for these analyses, which pose significant challenges to their implementation in clinical diagnostic laboratories. Here, we describe, to our knowledge, the first evaluation of various processing techniques and cryopreservation methods combined with a new designed spectral flow cytometry panel for high dimensional immunophenotyping. This study highlights both limitations and possibilities for clinical implementation of immunophenotyping of blood liquid biopsies.

The design and employment of a single 37-marker SFC designed for the evaluation of circulating immune cell composition, differentiation, and activation status within a blood liquid biopsy was essential to our work. By incorporating a wide range of markers into this one panel we have established an unbiased, in-depth analysis of immune phenotypes beyond classical standards which we believe have clinical relevance. This HD-ICPP assay coupled with stringent gating strategies and data visualization gives an easy and reliable roadmap to implement it in clinical diagnostics laboratories.

A major objective of our study was to investigate if a simple, rapid, and cost-effective sampling and processing approach could become applicable to diagnostic laboratories. This combined with the 37-marker SFC panel gives, for the first time, a very detailed interpretation of how blood liquid biopsies may be handled to limit off-target data analysis effects. Overall, any cryopreservation method, based on limited processing of a blood sample, was associated with a better comparison to the fresh blood samples. We found that SepMate-Ficoll gradient purification performed well but considering the need for at least 2 hours of processing before cryopreservation this method may be too time-consuming for a clinical diagnostic setting. The faster HepaSep purification approach did not perform any better that SepMate-Ficoll, and thus we argue that the rapid handling of merely cryopreserve whole blood outperformed any of the other methods. The usage of RBC lysis was rapid and seemed not to induce any confounding effects on activation or differentiation markers within the cryopreserved samples.

Many different cryopreservation matrices exist, with the majority being based on the solvent DMSO. We chose to test three methods, two commercially available matrices and a classical research-grade created protocol. With our HC-ICPP assay, we had the possibility to conduct detailed performance analysis, including overall cell composition as well as rare and specific cellular subset. Considering all the various analyses conducted across our 6 donors, CS10 performed best based on preserving the phenotypic signature, though all methods had some impact on expression markers, especially CD28. Such observations raise some concerns for past and future work on immunophenotypic analysis. We argue that there is a general need to conduct validation of phenotypic markers comparing signals from fresh and frozen samples, thereby avoiding inappropriate data interpretations.

In conclusion, our work adds important insight into the feasibility of implementing a tool as the HD-ICPP in clinical settings to support monitoring therapy effects and host immune competence over disease development. We demonstrate a new and cancer-relevant optimized 37-marker flow protocol applicable for assessing immune cell distribution and function, and believe this offers solutions to the field on how easy processing of liquid biopsies can be achieved - that actually reflects the condition of a patient at the time the blood sample has been collected.

## Methods

### Spectral flow panel: Design, optimization and validation

A 37-marker spectral flow panel for comprehensive immunophenotyping of human peripheral blood mononuclear cells (PBMCs) was designed and validated on a SONY ID7000™ Spectral Cell Analyzer equipped with 5 lasers (355, 405, 488, 561, and 637 nm) and 147 fluorescence detectors (SONY Biotechnologies, San Jose, CA). First, markers needed to identify and characterize immune cell populations were selected. Next, markers were matched with fluorochromes while considering the expression level of the markers and the fluorescence intensity of the fluorochromes so that lowly expressed markers were matched with bright fluorochromes and vice versa. At the same time, the combination of fluorochromes was theoretically assessed by the quantification of a similarity matrix on the SONY ID7000 Spectral Panel Design Tool (Figure S2). Here, a spillover similarity index between 0 and 1 was calculated for multiple pairs of fluorochromes. A high index indicates a high degree of spectral similarity and, hence, fluorochromes with a high index were excluded or assigned to antigens that are mutually exclusively expressed on PBMCs to prevent data spreading.

Next, titration experiments were carried out on frozen PBMCs to determine the optimal antibody concentrations. All antibodies were tested in a 2-fold serial dilution in Fluorescence-activated cell sorting (FACS) wash buffer (Phosphate buffered saline (PBS, BioWest, cat# L0615-500) + 0.5% Bovine serum albumin (BSA, Sigma-Aldrich, cat# A8806-5G) + 0.09% Na-azide (Sigma-Aldrich, cat# S2002-5G + 1 mM Ethylenediaminetetraacetic acid (EDTA, Invitrogen, cat# 15575020)). Titration performance was then assessed in FlowJO™ version 10.9.0 by calculating the stain index and evaluating the separation of the negative and positive population for each marker. For the titration of activation- and checkpoint markers, thawed PBMCs were activated prior to staining with either aCD3 (eBioscience, cat# 16-0037-85)/CD28 (eBioscience, cat# 16-0289-85), IFNg (PeproTech, cat# 300-02L), or PMA (Sigma-Aldrich, cat# 79346-5MG) + ionomycin (Sigma-Aldrich, cat# I9657-1MG) for 24hrs to further estimate the expression level of markers through histograms. Lower antibody concentrations than those suggested by the stain index were selected if the separation between negative and positive population was considered good. For antibodies with poor titration performance, new clones and fluorochromes were tested and titrated.

Finally, a multicolor optimization was conducted on both activated (PMA + ionomycin) and nonactivated frozen PBMCs. Individual marker performance was then assessed by comparing multicolored samples and single stained samples. Extensive data spread into channels was evaluated, and fluorochromes with a high degree of data spread were replaced to improve the resolution of markers in the panel. Different combinations of master mixes, as well as Fluorescence Minus One (FMO)s for different activation- and checkpoint markers, were evaluated. Based on the abovementioned assessments, a 37-marker spectral flow panel was developed for the characterization of human PBMCs, including lineage, differentiation, activation and checkpoint markers (see Table 1).

### Blood samples and experimental setup

Whole blood samples were collected as buffy coats obtained from six anonymized healthy adult blood donors from Aarhus University Hospital (AUH) Blood Bank. Studies on anonymized samples are exempt from ethical permissions in Denmark (Kommiteeloven §14 stk. 3). Undiluted whole blood (WB) was divided into three different tubes based on the isolation method (WB, HetaSep, Ficoll). For each isolation method, blood samples were divided into four additional tubes and either analyzed fresh by flow cytometry or cryopreserved prior to flow cytometry analysis using three different cryopreservation mediums – BloodStor® 55-5 (BS55-5, STEMCELL, cat# 07937), CryoStor10 (CS10, STEMCELL, cat# 07930) or standard freezing medium (SFM, 90% fetal calf serum (FCS) and 10% Dimethylsulfoxid (DMSO)) (See Figure S5).

### PBMC isolation

#### Whole blood

For WB, cells underwent RBC lysis (ACK lysis buffer, Gibco, cat# A10492-01) prior to further processing, except for cells undergoing BS55-5-cryopreservation. Subsequently, cells were centrifuged (300 x g for 5 min) and counted. A total of 5 × 10^6^ cells were taken out for spectral flow staining, while the rest of the cells were cryopreserved.

#### HetaSep

One third of the undiluted WB was isolated with HetaSep™ (STEMCELL, cat# 07906) according to the manufacturer’s instructions. One part of HetaSep™ was added to five parts of whole blood and the sample was allowed to settle until plasma and RBC layer were separated sufficiently. The leukocyte-rich plasma layer was harvested and placed in another tube and washed in 4-fold complete RPMI (cRPMI) (RPMI (Sigma-Aldrich, cat# R8758-500ML) + 10% FCS + 1% L-Glu + 1% P/S). Next, cells were centrifuged at 120 x g for 10 minutes with the brake off. Supernatant was discarded and cells were resuspended in cRPMI and counted. 5 × 10^6^ cells were used for spectral flow staining, while the rest of the sample was equally divided into three different tubes based on cryopreservation method.

#### Ficoll

For the isolation of PBMCs with Ficoll, 13 mL of Ficoll-Paque PLUS (GE Healthcare, cat# 17-1440-03) was added to the insert of a SepMate tube (STEMCELL, cat# 85460). Whole blood was diluted by adding an equal volume of PBS. Hereafter, the diluted whole blood was gently added to the SepMate tube without the blood getting into the insert. The SepMate tube was then centrifuged at 1200 x g for 10 minutes with the brake on and the leukocyte-rich plasma was then poured into a new tube. Cells were washed in PBS and centrifuged with the brake on. Subsequently, cells were counted, and 5 × 10^6^ cells were used for spectral flow and the rest were cryopreserved.

### Cryopreservation of PBMCs

#### BloodStor® 55-5

Five parts of undiluted whole blood was mixed with one part of BloodStor® 55-5 (BS55-5, STEMCELL, cat# 07937) and cryopreserved. HetaSep- and Ficoll-isolated PBMCs were first resuspended in FCS for a concentration of 10 mio cells/ml and one fifth of BS55-5 was added to the cell suspension and cryopreserved.

#### Standard freezing medium

RBC lysed whole blood cells as well as HetaSep- and Ficoll-isolated PBMCs were resuspended in SFM (90% FCS (Sigma-Aldrich, cat# F9665) and 10% DMSO (Sigma-Aldrich, cat# D4540) for a concentration of 10 mio cells/ml and cryopreserved.

#### CryoStor10

RBC lysed whole blood cells, HetaSep- and Ficoll-isolated PBMCs were resuspended in CryoStor10 (STEMCELL, cat# 07930) for a concentration of 10 mio cells/ml. As described by the manufacturer, the cell suspension was incubated for approximately 10 minutes at 2-8°C before freezing.

All cryopreserved cells were transferred to a −80°C freezer in cell freezing containers for at least 12 hours and, afterwards, stored at −150°C until flow analysis (approx. after 7 days).

### Thawing

Cryopreserved samples were thawed by carefully resuspending the cells in preheated (37°C) cRPMI and the cell suspension was transferred to a 15 ml conical tube. Cells were then centrifuged at 300 x g for 4 min and resuspended in cRPMI. In addition, BloodStor-cryopreserved whole blood samples underwent RBC lysis before spectral flow staining. Thawed cells were then transferred to a 96-well plate.

### Staining procedure

Generally, all staining procedures were done at room temperature and protected from sunlight. First, PBMCs were washed in PBS and then stained with LiveDead FixableBlue viability dye (Thermo Scientific, cat# L34961) (1:100 dilution in PBS) for 15 min. Subsequently, cells were washed by adding 200 µl PBS and centrifuged at 300 x g for 4 minutes. Next, cells were blocked using human IgG (100 µg/ml) (Sigma Aldrich, cat# I2511-10MG) for 30 minutes. After incubation, 50 µl of Mastermix 1 (Table S1) was added to each of the multicolor samples and allowed to incubate for 30 minutes. Mastermix 2 (antibodies diluted in TrueStain Monocyte Blocker™ (BioLegend, cat# 426103), CellBlox Blocking Buffer (Thermo Fisher Scientific, cat# C001T06F01), and FACS wash (PBS + 0.5% BSA (Bovine serum albumin) + 0.09% Na-azide + 1 mM EDTA (ethylenediaminetetraacetic acid))) was subsequently added to the multicolor samples, and cells were incubated for 30 minutes. Next, cells were washed by adding 100 µl FACS wash and centrifuged at 300 x g for 4 minutes. Subsequently, cells were resuspended in 50 µl IgG (100 ug/ml) followed by 100 µl Antibody Mastermix 3+4 (including Brilliant Stain Buffer Plus (BD bioscience, cat# 566385), TrueStain Monocyte Blocker and FACS wash) and incubated for 60 minutes. Afterwards, the cells were washed by adding 200 µl FACS wash and centrifuged at 300 x g for 4 minutes and then fixated and permeabilized using the FoxP3/Transcription Factor staining buffer set (eBioscience, cat# 00-5523-00) prior to intracellular staining. Cells were incubated in FoxP3 fixation/permeabilization working solution (1 part of Foxp3 Fixation/Permeabilization Concentrate to 3 parts of Foxp3 Fixation/Permeabilization Diluent) for 30 minutes. Afterwards, cells were washed twice with 1X permeabilization buffer (1 part of 10X Permeabilization Buffer with 9 parts of distilled water) (500 x g for 5 min) and blocked with IgG (100 µg/ml) diluted in 1X permeabilization buffer for 10 minutes. A total of 50 µl of Mastermix 5 (including Brilliant Stain Buffer Plus and 1X Permeabilization buffer) was then added, and cells were incubated for 30 minutes. Cells were washed two times by adding 200 µl 1X permeabilization buffer and pelleted by centrifugation at 500 x g for 5 min. Finally, cells were resuspended in 100 µl FACS wash before running the samples on the SONY ID7000™ Spectral Cell Analyzer.

Both unstained control samples per donor and condition as well as single stained controls beads were included and processed following the same procedure as the multicolored samples for the spectral unmixing. UltraComp eBeads™ Plus Compensation Beads (Invitrogen, cat# 01-3333-42) were used for the extracellular and intracellular antibodies, while ArC™ Amine Reactive Compensation Beads (Life Technologies, cat# A10346) were used for the LiveDead FixableBlue viability dye.

### Data acquisition and unmixing

Samples were run on the SONY ID7000™ Spectral Cell Analyzer on normalized mode, and data was acquired and unmixed on the appertaining SONY ID7000™ Acquisition and Analysis Software version 2.0.2 (SONY Biotechnologies, San Jose, CA) utilizing the Weighted Least Squares Method (WLSM) algorithm for spectral unmixing. Unmixing was calculated based on the negative and positive spectrum from the included single stained control beads and the autofluorescence exhibited by the included unstained controls. The calculated unmixing matrix was verified by screening screening NxN pairwise plots plotting all channels against each other. The verified unmixing matrix was then applied to all samples.

### Data analysis and statistics

Unmixed Flow cytometry standard (FCS) files were imported to FlowJo™ (v10) and CytoBank Premium (Beckman Coulter, Inc.) and analyzed. Manual gating was performed based on established gating strategies and immune cell frequencies were extracted. Statistical analysis was performed in GraphPad Prism version 10. One-way ANOVA analysis with Bonferroni’s multiple comparison test was used to compare frequencies and MFIs for each immune cell type between isolation- and cryopreservation methods. Pearson’s correlation coefficient test was conducted to compare immune cell frequencies and MFI of functional markers between whole blood and different isolation and cryopreservation methods. Unsupervised dimensionality reduction analyses based on Uniform Manifold Approximation and Projection (UMAP) and t-distributed stochastic neighbor embedding (tSNE) were run on manually gated live CD45^+^ singlets from one representative donor, live CD3^+^CD4^+^ and CD3^+^CD8^+^ singlets, and live CD3^+^CD14^-^ and CD3^-^CD14^+^ singlets. UMAP settings were as follows: Input parameters for the live CD45^+^ singlets CD4, CD3, CD16, HLA-DR, CD11c, CD14, CD123, CD8, CD19, CD66b, and CD56, input parameters for live CD3^+^CD14^-^ and CD3^-^CD14^+^ singlets Forward Scatter (FSC), Side Scatter (SSC), CD3, CD14, CD4, CD8, and HLA-DR, Distance Function = Euclidian, Nearest Neighbors = 15, Minimum distance = 0.5, # of components = 2. tSNE settings were as follows: Input parameters CD27, CD45RA, and CCR7, Iterations = 1000, Perplexity = 30, Learning rate = 14,000 (live CD3^+^CD4^+^ singlets) and 12,324 (live CD3^+^CD8^+^ singlets). All samples were downsized to 50,000 events and concatenated. Furthermore, unsupervised clustering was carried out based on the FlowSOM algorithm in FlowJo™. FlowSOM settings were as follows: Input parameters for live CD45+ singlets CD4, CD3, CD16, HLA-DR, CD11c, CD14, CD123, CD8, CD19, CD66b, and CD56, input parameters for live CD3^+^CD14^-^ and CD3^-^CD14^+^ singlets Forward Scatter (FSC), Side Scatter (SSC), CD3, CD14, CD4, CD8, and HLA-DR, input parameters for live CD3^+^CD4^+^ and CD3^+^CD8^+^ singlets CD27, CD45RA, and CCR7, Number of meta clusters = 9 (live CD45^+^ singlets, figure 1B), 10 (live CD45^+^ singlets, figure 3A), 7 (live CD3^+^CD4^+^ and CD3^+^CD8^+^ singlets), and 3 (live CD3^+^CD14^-^ and CD3^-^CD14^+^ singlets), SOM grid size (W x H) = 10 × 10, Node scale = 100%.

## Acknowledgement and Funding

This work was supported and funded by The Danish Cancer Society (R246-A14695), Novo Nordisk Foundation (NNF20OC0062825), Danish Kræftforsknings foundation, Riisfort Foundation, and Dagmar Marshalls Foundation.

The authors would like to thank the FACS Core facility at Aarhus University for their technical and methodological support, as well as for their maintenance and service of the ID7000, which ensured stable and proper machine function. The 5-laser ID7000 is a generous gift from the Novo Nordisk Foundation, grant number NNF210C0066798. Additionally, we would like to thank the FACS core facility for providing valuable feedback on the technical aspects of the manuscript.

The authors would also like to thank João Monteiro from SONY Biotechnologies for his assistance in the initial design of the panel, as well as for his ongoing support on technical aspects of operating the ID7000 and performing correct unmixing in the SONY analyzer software.

## Author contribution

Conceptualization: JGP, MRJ; Methodology: JGP, JJCL, ET, KRG; Investigation/ acquisition of data: JGP, JJCL, ET, LLD, KGR; Data analysis: JGP, JJCL, MRJ; Writing and reviewing; JGP, JJCL, ET, LLD, AWL, KRG, and MRJ; Visualization: JGP, JJCL and MRJ; Study supervision: MRJ.

