## Supplemental figures for "Using high-dimensional immunophenotyping for mapping immune cell composition in rapid cryopreserved liquid biopsies"

#### SUPPLEMENTARY FIGURES

Pedersen et al. figure S1

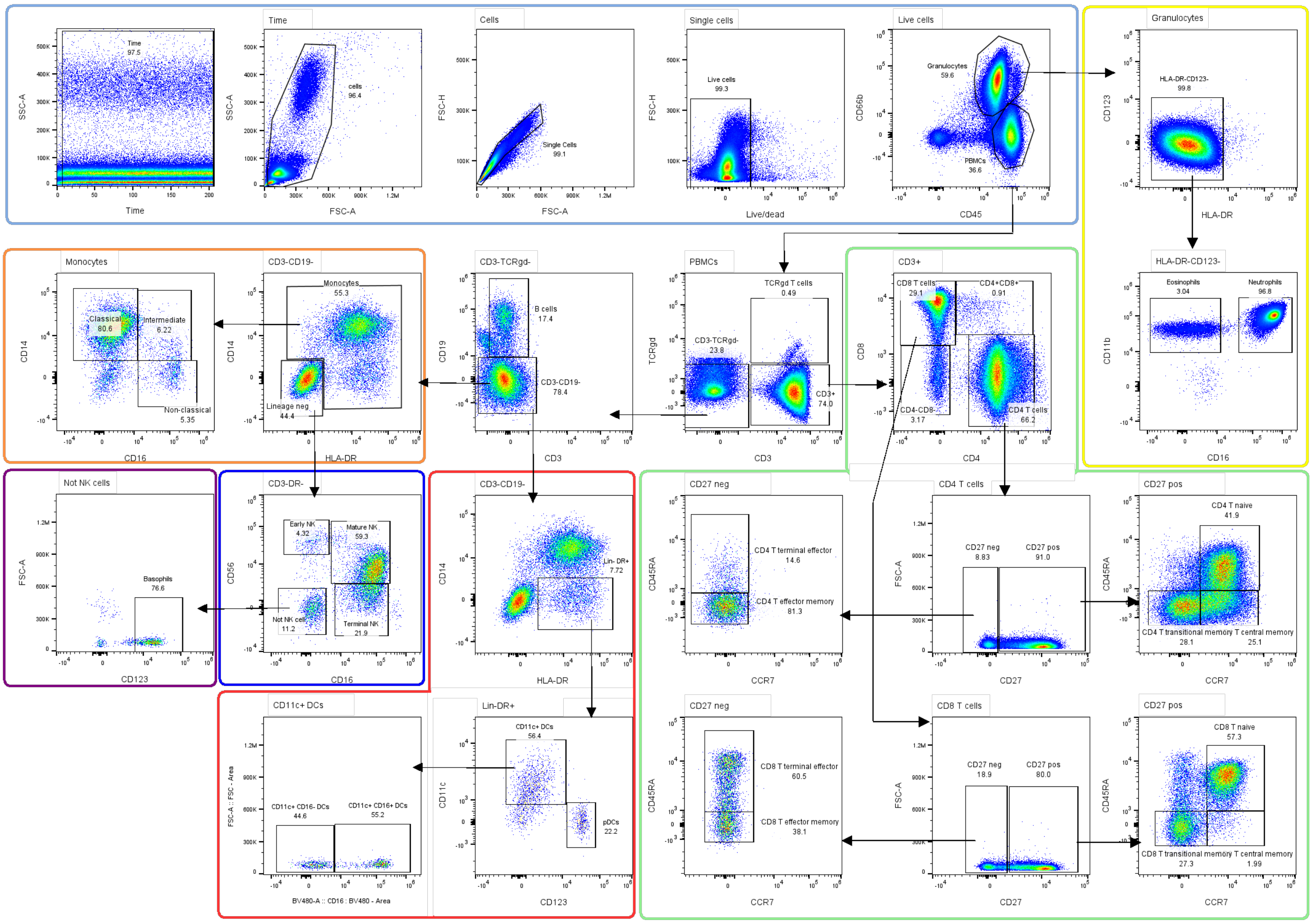

**Figure S1. The manual gating strategy used to define each cell population.** Representative example of one fresh blood sample stained with the 37-marker panel, focusing on manual gating of all the major subpopulations within the CD45+ compartment. Population frequencies represent the frequency of parent population. Gating was done based on FMO and single stained controls.

### Pedersen et al. figure S2

| | | | CD73 | Live/dead | CD28 | CD69 | TIM-3 | CD25 | PD-1 | IFNg | STING | CD45 | CD16 | HLA-DR | CD123 | CD27 | CD86 | CD163 | CTLA-4 |
|---|---|---|---|---|---|---|---|---|---|---|---|---|---|---|---|---|---|---|---|
| Marker | Fluorochrome | Sum | BUV395 | LiveDeadFixableBlue | BUV496 | BUV563 | BUV615 | BUV661 | BUV737 | BUV805 | BV421 | PacBlue | BV480 | BV570 | BV605 | BV650 | BV711 | BV750 | BV785 |
| CD73 | BUV395 | 0.4 | | 0.17 | 0.02 | 0.01 | 0.00 | 0.00 | 0.01 | 0.15 | 0.01 | 0.00 | 0.00 | 0.00 | 0.00 | 0.00 | 0.00 | 0.00 | 0.00 |
| Live/dead | LiveDeadFixableBlue | 0.8 | 0.17 | | 0.22 | 0.01 | 0.00 | 0.00 | 0.00 | 0.05 | 0.10 | 0.02 | 0.02 | 0.00 | 0.00 | 0.01 | 0.01 | 0.00 | 0.00 |
| CD28 | BUV496 | 0.6 | 0.02 | 0.22 | | 0.03 | 0.00 | 0.01 | 0.00 | 0.00 | 0.01 | 0.01 | 0.14 | 0.00 | 0.00 | 0.01 | 0.01 | 0.01 | 0.00 |
| CD69 | BUV563 | 1.0 | 0.01 | 0.01 | 0.03 | | 0.24 | 0.00 | 0.01 | 0.00 | 0.00 | 0.00 | 0.00 | 0.17 | 0.06 | 0.01 | 0.03 | 0.03 | 0.02 |
| TIM-3 | BUV615 | 1.0 | 0.00 | 0.00 | 0.00 | 0.24 | | 0.03 | 0.01 | 0.00 | 0.00 | 0.01 | 0.01 | 0.08 | 0.27 | 0.01 | 0.02 | 0.02 | 0.02 |
| CD25 | BUV661 | 1.4 | 0.00 | 0.00 | 0.01 | 0.00 | 0.03 | | 0.01 | 0.00 | 0.00 | 0.01 | 0.01 | 0.02 | 0.00 | 0.15 | 0.02 | 0.01 | 0.02 |
| PD-1 | BUV737 | 0.9 | 0.01 | 0.00 | 0.00 | 0.01 | 0.01 | 0.01 | | 0.05 | 0.00 | 0.01 | 0.01 | 0.03 | 0.02 | 0.00 | 0.16 | 0.13 | 0.02 |
| IFNg | BUV805 | 0.6 | 0.15 | 0.05 | 0.00 | 0.00 | 0.00 | 0.00 | 0.05 | | 0.00 | 0.00 | 0.00 | 0.02 | 0.02 | 0.02 | 0.00 | 0.01 | 0.07 |
| STING | BV421 | 0.9 | 0.01 | 0.10 | 0.01 | 0.00 | 0.00 | 0.00 | 0.00 | 0.00 | | 0.43 | 0.05 | 0.01 | 0.00 | 0.00 | 0.00 | 0.01 | 0.12 |
| CD45 | PacBlue | 1.2 | 0.00 | 0.02 | 0.01 | 0.00 | 0.01 | 0.01 | 0.01 | 0.00 | 0.43 | | 0.46 | 0.00 | 0.00 | 0.00 | 0.00 | 0.00 | 0.05 |
| CD16 | BV480 | 0.9 | 0.00 | 0.02 | 0.14 | 0.00 | 0.01 | 0.01 | 0.01 | 0.00 | 0.05 | 0.46 | | 0.00 | 0.00 | 0.01 | 0.01 | 0.00 | 0.00 |
| HLA-DR | BV570 | 1.9 | 0.00 | 0.00 | 0.00 | 0.17 | 0.08 | 0.02 | 0.03 | 0.02 | 0.01 | 0.00 | 0.00 | | 0.48 | 0.01 | 0.03 | 0.03 | 0.01 |
| CD123 | BV605 | 1.5 | 0.00 | 0.00 | 0.00 | 0.06 | 0.27 | 0.00 | 0.02 | 0.02 | 0.00 | 0.00 | 0.00 | 0.48 | | 0.14 | 0.01 | 0.02 | 0.02 |
| CD27 | BV650 | 0.9 | 0.00 | 0.01 | 0.01 | 0.01 | 0.01 | 0.15 | 0.00 | 0.02 | 0.00 | 0.00 | 0.01 | 0.01 | 0.14 | | 0.04 | 0.00 | 0.01 |
| CD86 | BV711 | 1.7 | 0.00 | 0.01 | 0.01 | 0.03 | 0.02 | 0.02 | 0.16 | 0.00 | 0.00 | 0.00 | 0.01 | 0.03 | 0.01 | 0.04 | | 0.19 | 0.03 |
| CD163 | BV750 | 1.4 | 0.00 | 0.00 | 0.01 | 0.03 | 0.02 | 0.01 | 0.13 | 0.01 | 0.01 | 0.00 | 0.00 | 0.03 | 0.02 | 0.00 | 0.19 | | 0.51 |
| CTLA-4 | BV785 | 1.3 | 0.00 | 0.00 | 0.00 | 0.02 | 0.02 | 0.02 | 0.02 | 0.07 | 0.12 | 0.05 | 0.00 | 0.01 | 0.02 | 0.01 | 0.03 | 0.51 | |
| CD19 | BB515 | 1.3 | 0.00 | 0.00 | 0.00 | 0.00 | 0.01 | 0.01 | 0.01 | 0.00 | 0.00 | 0.00 | 0.00 | 0.01 | 0.01 | 0.01 | 0.01 | 0.01 | 0.01 |
| CD66b | FITC | 1.4 | 0.00 | 0.00 | 0.00 | 0.00 | 0.01 | 0.01 | 0.01 | 0.00 | 0.00 | 0.00 | 0.00 | 0.01 | 0.01 | 0.01 | 0.02 | 0.01 | 0.01 |
| CD14 | NovaFluorBlue585 | 1.6 | 0.00 | 0.01 | 0.00 | 0.00 | 0.00 | 0.02 | 0.02 | 0.01 | 0.00 | 0.01 | 0.01 | 0.00 | 0.00 | 0.02 | 0.03 | 0.02 | 0.03 |
| CD4 | NovaFluorBlue610 | 1.7 | 0.00 | 0.00 | 0.00 | 0.00 | 0.01 | 0.01 | 0.02 | 0.01 | 0.00 | 0.01 | 0.01 | 0.01 | 0.01 | 0.01 | 0.03 | 0.03 | 0.03 |
| CD8 | PerCP | 1.0 | 0.00 | 0.01 | 0.01 | 0.01 | 0.00 | 0.08 | 0.00 | 0.01 | 0.01 | 0.01 | 0.02 | 0.01 | 0.00 | 0.14 | 0.00 | 0.01 | 0.02 |
| CD11c | BB700 | 1.4 | 0.01 | 0.02 | 0.01 | 0.02 | 0.02 | 0.00 | 0.00 | 0.03 | 0.01 | 0.01 | 0.01 | 0.02 | 0.01 | 0.01 | 0.07 | 0.00 | 0.03 |
| TCRgd | PerCP-eFluor710 | 2.0 | 0.01 | 0.02 | 0.03 | 0.05 | 0.03 | 0.00 | 0.03 | 0.02 | 0.02 | 0.03 | 0.03 | 0.06 | 0.03 | 0.00 | 0.35 | 0.03 | 0.00 |
| CD38 (SBB765) | BB755 | 0.8 | 0.00 | 0.01 | 0.01 | 0.02 | 0.02 | 0.01 | 0.05 | 0.00 | 0.01 | 0.01 | 0.01 | 0.03 | 0.03 | 0.02 | 0.01 | 0.14 | 0.05 |
| FOXP3 | PE | 2.2 | 0.00 | 0.00 | 0.00 | 0.10 | 0.01 | 0.01 | 0.01 | 0.01 | 0.00 | 0.00 | 0.01 | 0.29 | 0.02 | 0.01 | 0.02 | 0.02 | 0.02 |
| CD56 | SparkYG-581 | 2.0 | 0.00 | 0.00 | 0.00 | 0.08 | 0.04 | 0.01 | 0.01 | 0.01 | 0.00 | 0.00 | 0.00 | 0.34 | 0.07 | 0.01 | 0.02 | 0.01 | 0.01 |
| CCR7 | PE-Dazzle594 | 2.2 | 0.00 | 0.01 | 0.01 | 0.02 | 0.12 | 0.01 | 0.02 | 0.01 | 0.01 | 0.01 | 0.01 | 0.09 | 0.09 | 0.01 | 0.04 | 0.03 | 0.03 |
| CD3 | PE-Fire640 | 1.0 | 0.00 | 0.01 | 0.01 | 0.01 | 0.00 | 0.01 | 0.02 | 0.02 | 0.01 | 0.02 | 0.02 | 0.00 | 0.00 | 0.00 | 0.03 | 0.04 | 0.05 |
| ICOS | PE-Cy5.5 | 1.2 | 0.00 | 0.01 | 0.01 | 0.01 | 0.00 | 0.01 | 0.02 | 0.02 | 0.01 | 0.01 | 0.02 | 0.00 | 0.01 | 0.01 | 0.01 | 0.03 | 0.04 |
| Ki67 | PE-Vio770 | 0.6 | 0.00 | 0.01 | 0.00 | 0.01 | 0.01 | 0.02 | 0.01 | 0.00 | 0.00 | 0.00 | 0.01 | 0.02 | 0.03 | 0.03 | 0.03 | 0.01 | 0.00 |
| PD-L1 | PE-Fire810 | 1.4 | 0.01 | 0.01 | 0.02 | 0.01 | 0.00 | 0.04 | 0.03 | 0.01 | 0.01 | 0.02 | 0.02 | 0.05 | 0.00 | 0.05 | 0.06 | 0.03 | 0.03 |
| Tbet | APC | 1.8 | 0.00 | 0.00 | 0.01 | 0.01 | 0.00 | 0.43 | 0.00 | 0.01 | 0.00 | 0.01 | 0.01 | 0.01 | 0.01 | 0.07 | 0.00 | 0.01 | 0.01 |
| CD11b | AF647 | 2.0 | 0.00 | 0.00 | 0.00 | 0.01 | 0.01 | 0.33 | 0.00 | 0.01 | 0.00 | 0.01 | 0.01 | 0.02 | 0.01 | 0.02 | 0.02 | 0.01 | 0.01 |
| TIGIT | AF660 | 1.9 | 0.00 | 0.01 | 0.01 | 0.02 | 0.01 | 0.10 | 0.01 | 0.01 | 0.01 | 0.01 | 0.01 | 0.03 | 0.02 | 0.00 | 0.10 | 0.01 | 0.02 |
| CD45RA | AF700 | 1.4 | 0.00 | 0.01 | 0.01 | 0.02 | 0.01 | 0.02 | 0.11 | 0.00 | 0.00 | 0.01 | 0.01 | 0.02 | 0.02 | 0.00 | 0.28 | 0.00 | 0.00 |
| LAG-3 | APC-Cy7 | 0.4 | 0.00 | 0.01 | 0.01 | 0.02 | 0.01 | 0.00 | 0.01 | 0.01 | 0.00 | 0.01 | 0.01 | 0.02 | 0.02 | 0.01 | 0.01 | 0.01 | 0.01 |

| | | CD19 | CD66b | CD14 | CD4 | CD8 | CD11c | TCRgd | CD38 (SBB765 | FOXP3 | CD56 | CCR7 | CD3 | ICOS | Ki67 | PD-L1 | Tbet | CD11b | TIGIT | CD45RA | LAG-3 |
|---|---|---|---|---|---|---|---|---|---|---|---|---|---|---|---|---|---|---|---|---|---|
| Marker | Fluorochrome | BB515 | FITC | NovaFluorBlue585 | NovaFluorBlue610 | PerCP | BB700 | PerCP-eFluor710 | BB755 | PE | SparkYG-581 | PE-Dazzle594 | PE-Fire640 | PE-Cy5.5 | PE-Vio770 | PE-Fire810 | APC | AF647 | AF660 | AF700 | APC-Cy7 |
| CD73 | BUV395 | 0.00 | 0.00 | 0.00 | 0.00 | 0.00 | 0.01 | 0.01 | 0.00 | 0.00 | 0.00 | 0.00 | 0.00 | 0.00 | 0.00 | 0.01 | 0.00 | 0.00 | 0.00 | 0.00 | 0.00 |
| Live/dead | LiveDeadFixableBlue | 0.00 | 0.00 | 0.01 | 0.00 | 0.01 | 0.02 | 0.02 | 0.01 | 0.00 | 0.00 | 0.01 | 0.01 | 0.01 | 0.01 | 0.01 | 0.00 | 0.00 | 0.01 | 0.01 | 0.01 |
| CD28 | BUV496 | 0.00 | 0.00 | 0.00 | 0.00 | 0.01 | 0.01 | 0.03 | 0.01 | 0.00 | 0.00 | 0.01 | 0.01 | 0.01 | 0.00 | 0.02 | 0.01 | 0.00 | 0.01 | 0.01 | 0.01 |
| CD69 | BUV563 | 0.00 | 0.00 | 0.00 | 0.00 | 0.01 | 0.02 | 0.05 | 0.02 | 0.10 | 0.08 | 0.02 | 0.01 | 0.01 | 0.01 | 0.01 | 0.01 | 0.01 | 0.02 | 0.02 | 0.02 |
| TIM-3 | BUV615 | 0.01 | 0.01 | 0.00 | 0.01 | 0.00 | 0.02 | 0.03 | 0.02 | 0.01 | 0.04 | 0.12 | 0.00 | 0.00 | 0.01 | 0.00 | 0.00 | 0.01 | 0.01 | 0.01 | 0.01 |
| CD25 | BUV661 | 0.01 | 0.01 | 0.02 | 0.01 | 0.08 | 0.00 | 0.00 | 0.01 | 0.01 | 0.01 | 0.01 | 0.01 | 0.01 | 0.02 | 0.04 | 0.43 | 0.33 | 0.10 | 0.02 | 0.00 |
| PD-1 | BUV737 | 0.01 | 0.01 | 0.02 | 0.02 | 0.00 | 0.00 | 0.03 | 0.05 | 0.01 | 0.01 | 0.02 | 0.02 | 0.02 | 0.01 | 0.03 | 0.00 | 0.00 | 0.01 | 0.11 | 0.01 |
| IFNg | BUV805 | 0.00 | 0.00 | 0.01 | 0.01 | 0.01 | 0.03 | 0.02 | 0.00 | 0.01 | 0.01 | 0.01 | 0.02 | 0.02 | 0.00 | 0.01 | 0.01 | 0.01 | 0.01 | 0.00 | 0.01 |
| STING | BV421 | 0.00 | 0.00 | 0.00 | 0.00 | 0.01 | 0.01 | 0.02 | 0.01 | 0.00 | 0.00 | 0.01 | 0.01 | 0.01 | 0.00 | 0.01 | 0.00 | 0.00 | 0.01 | 0.00 | 0.00 |
| CD45 | PacBlue | 0.00 | 0.00 | 0.01 | 0.01 | 0.01 | 0.01 | 0.03 | 0.01 | 0.00 | 0.00 | 0.01 | 0.02 | 0.01 | 0.00 | 0.02 | 0.01 | 0.01 | 0.01 | 0.01 | 0.01 |
| CD16 | BV480 | 0.00 | 0.00 | 0.01 | 0.01 | 0.02 | 0.01 | 0.03 | 0.01 | 0.01 | 0.00 | 0.01 | 0.02 | 0.02 | 0.01 | 0.02 | 0.01 | 0.01 | 0.01 | 0.01 | 0.01 |
| HLA-DR | BV570 | 0.01 | 0.01 | 0.00 | 0.01 | 0.01 | 0.02 | 0.06 | 0.03 | 0.29 | 0.34 | 0.09 | 0.00 | 0.00 | 0.02 | 0.05 | 0.01 | 0.02 | 0.03 | 0.02 | 0.02 |
| CD123 | BV605 | 0.01 | 0.01 | 0.00 | 0.01 | 0.00 | 0.01 | 0.03 | 0.03 | 0.02 | 0.07 | 0.09 | 0.00 | 0.01 | 0.03 | 0.00 | 0.01 | 0.01 | 0.02 | 0.02 | 0.02 |
| CD27 | BV650 | 0.01 | 0.01 | 0.02 | 0.01 | 0.14 | 0.01 | 0.00 | 0.02 | 0.01 | 0.01 | 0.01 | 0.00 | 0.01 | 0.03 | 0.05 | 0.07 | 0.02 | 0.00 | 0.00 | 0.01 |
| CD86 | BV711 | 0.01 | 0.02 | 0.03 | 0.03 | 0.00 | 0.07 | 0.35 | 0.01 | 0.02 | 0.02 | 0.04 | 0.03 | 0.01 | 0.03 | 0.06 | 0.00 | 0.02 | 0.10 | 0.28 | 0.01 |
| CD163 | BV750 | 0.01 | 0.01 | 0.02 | 0.03 | 0.01 | 0.00 | 0.03 | 0.14 | 0.02 | 0.01 | 0.03 | 0.04 | 0.03 | 0.01 | 0.03 | 0.01 | 0.01 | 0.01 | 0.00 | 0.01 |
| CTLA-4 | BV785 | 0.01 | 0.01 | 0.03 | 0.03 | 0.02 | 0.03 | 0.00 | 0.05 | 0.02 | 0.01 | 0.03 | 0.05 | 0.04 | 0.00 | 0.03 | 0.01 | 0.01 | 0.02 | 0.00 | 0.01 |
| CD19 | BB515 | | 0.95 | 0.16 | 0.02 | 0.01 | 0.00 | 0.02 | 0.01 | 0.00 | 0.00 | 0.00 | 0.01 | 0.01 | 0.00 | 0.00 | 0.00 | 0.00 | 0.01 | 0.01 | 0.01 |
| CD66b | FITC | 0.95 | | 0.21 | 0.03 | 0.01 | 0.00 | 0.02 | 0.00 | 0.00 | 0.00 | 0.00 | 0.01 | 0.01 | 0.00 | 0.00 | 0.01 | 0.00 | 0.01 | 0.01 | 0.01 |
| CD14 | NovaFluorBlue585 | 0.16 | 0.21 | | 0.47 | 0.00 | 0.05 | 0.01 | 0.01 | 0.17 | 0.00 | 0.20 | 0.01 | 0.00 | 0.01 | 0.03 | 0.01 | 0.01 | 0.01 | 0.01 | 0.01 |
| CD4 | NovaFluorBlue610 | 0.02 | 0.03 | 0.47 | | 0.00 | 0.03 | 0.01 | 0.01 | 0.07 | 0.03 | 0.68 | 0.07 | 0.00 | 0.01 | 0.01 | 0.01 | 0.01 | 0.01 | 0.01 | 0.01 |
| CD8 | PerCP | 0.01 | 0.01 | 0.00 | 0.00 | | 0.23 | 0.05 | 0.00 | 0.01 | 0.00 | 0.00 | 0.12 | 0.09 | 0.02 | 0.02 | 0.01 | 0.01 | 0.03 | 0.00 | 0.01 |
| CD11c | BB700 | 0.00 | 0.00 | 0.05 | 0.03 | 0.23 | | 0.45 | 0.05 | 0.00 | 0.02 | 0.00 | 0.03 | 0.19 | 0.02 | 0.02 | 0.00 | 0.01 | 0.03 | 0.01 | 0.01 |
| TCRgd | PerCP-eFluor710 | 0.02 | 0.02 | 0.01 | 0.01 | 0.05 | 0.45 | | 0.19 | 0.03 | 0.02 | 0.02 | 0.01 | 0.28 | 0.00 | 0.02 | 0.00 | 0.00 | 0.08 | 0.10 | 0.00 |
| CD38 (SBB765) | BB755 | 0.01 | 0.00 | 0.01 | 0.01 | 0.00 | 0.05 | 0.19 | | 0.01 | 0.01 | 0.02 | 0.01 | 0.01 | 0.03 | 0.00 | 0.01 | 0.02 | 0.02 | 0.01 | 0.01 |
| FOXP3 | PE | 0.00 | 0.00 | 0.17 | 0.07 | 0.01 | 0.00 | 0.03 | 0.01 | | 0.70 | 0.23 | 0.00 | 0.02 | 0.00 | 0.34 | 0.01 | 0.01 | 0.01 | 0.01 | 0.01 |
| CD56 | SparkYG-581 | 0.00 | 0.00 | 0.00 | 0.03 | 0.00 | 0.02 | 0.02 | 0.01 | 0.70 | | 0.28 | 0.01 | 0.02 | 0.00 | 0.26 | 0.00 | 0.00 | 0.00 | 0.01 | 0.01 |
| CCR7 | PE-Dazzle594 | 0.00 | 0.00 | 0.20 | 0.68 | 0.00 | 0.00 | 0.02 | 0.02 | 0.23 | 0.28 | | 0.12 | 0.02 | 0.01 | 0.06 | 0.00 | 0.01 | 0.00 | 0.01 | 0.01 |
| CD3 | PE-Fire640 | 0.01 | 0.01 | 0.01 | 0.07 | 0.12 | 0.03 | 0.01 | 0.01 | 0.00 | 0.01 | 0.12 | | 0.21 | 0.01 | 0.00 | 0.05 | 0.02 | 0.05 | 0.00 | 0.01 |
| ICOS | PE-Cy5.5 | 0.01 | 0.01 | 0.00 | 0.00 | 0.09 | 0.19 | 0.28 | 0.01 | 0.02 | 0.02 | 0.02 | 0.21 | | 0.01 | 0.02 | 0.00 | 0.00 | 0.12 | 0.00 | 0.01 |
| Ki67 | PE-Vio770 | 0.00 | 0.00 | 0.01 | 0.01 | 0.02 | 0.02 | 0.00 | 0.03 | 0.00 | 0.00 | 0.01 | 0.01 | 0.01 | | 0.21 | 0.01 | 0.01 | 0.01 | 0.01 | 0.00 |
| PD-L1 | PE-Fire810 | 0.00 | 0.00 | 0.03 | 0.01 | 0.02 | 0.02 | 0.02 | 0.00 | 0.34 | 0.26 | 0.06 | 0.00 | 0.02 | 0.21 | | 0.02 | 0.02 | 0.02 | 0.02 | 0.00 |
| Tbet | APC | 0.00 | 0.01 | 0.01 | 0.01 | 0.01 | 0.00 | 0.00 | 0.01 | 0.01 | 0.00 | 0.00 | 0.05 | 0.00 | 0.01 | 0.02 | | 0.78 | 0.19 | 0.05 | 0.00 |
| CD11b | AF647 | 0.00 | 0.00 | 0.01 | 0.01 | 0.01 | 0.01 | 0.00 | 0.02 | 0.01 | 0.00 | 0.01 | 0.02 | 0.00 | 0.01 | 0.02 | 0.78 | | 0.47 | 0.12 | 0.01 |
| TIGIT | AF660 | 0.01 | 0.01 | 0.01 | 0.01 | 0.03 | 0.03 | 0.08 | 0.02 | 0.01 | 0.00 | 0.00 | 0.05 | 0.12 | 0.01 | 0.02 | 0.19 | 0.47 | | 0.43 | 0.01 |
| CD45RA | AF700 | 0.01 | 0.01 | 0.01 | 0.01 | 0.00 | 0.01 | 0.10 | 0.01 | 0.01 | 0.01 | 0.01 | 0.00 | 0.00 | 0.01 | 0.02 | 0.05 | 0.12 | 0.43 | | 0.08 |
| LAG-3 | APC-Cy7 | 0.01 | 0.01 | 0.01 | 0.01 | 0.01 | 0.01 | 0.00 | 0.01 | 0.01 | 0.01 | 0.01 | 0.01 | 0.01 | 0.00 | 0.00 | 0.00 | 0.01 | 0.01 | 0.08 | |

**Figure S2. Spillover spreading/similarity matrix.** Fluorochromes and related markers were paired and a spillover spreading score/similarity index between 0 and 1, representing the degree of spectral similarity. Fluorochromes with a high score/index were excluded or assigned to antigens that are mutually exclusively expressed on PBMCs to prevent data spreading.

Pedersen et al. figure S3

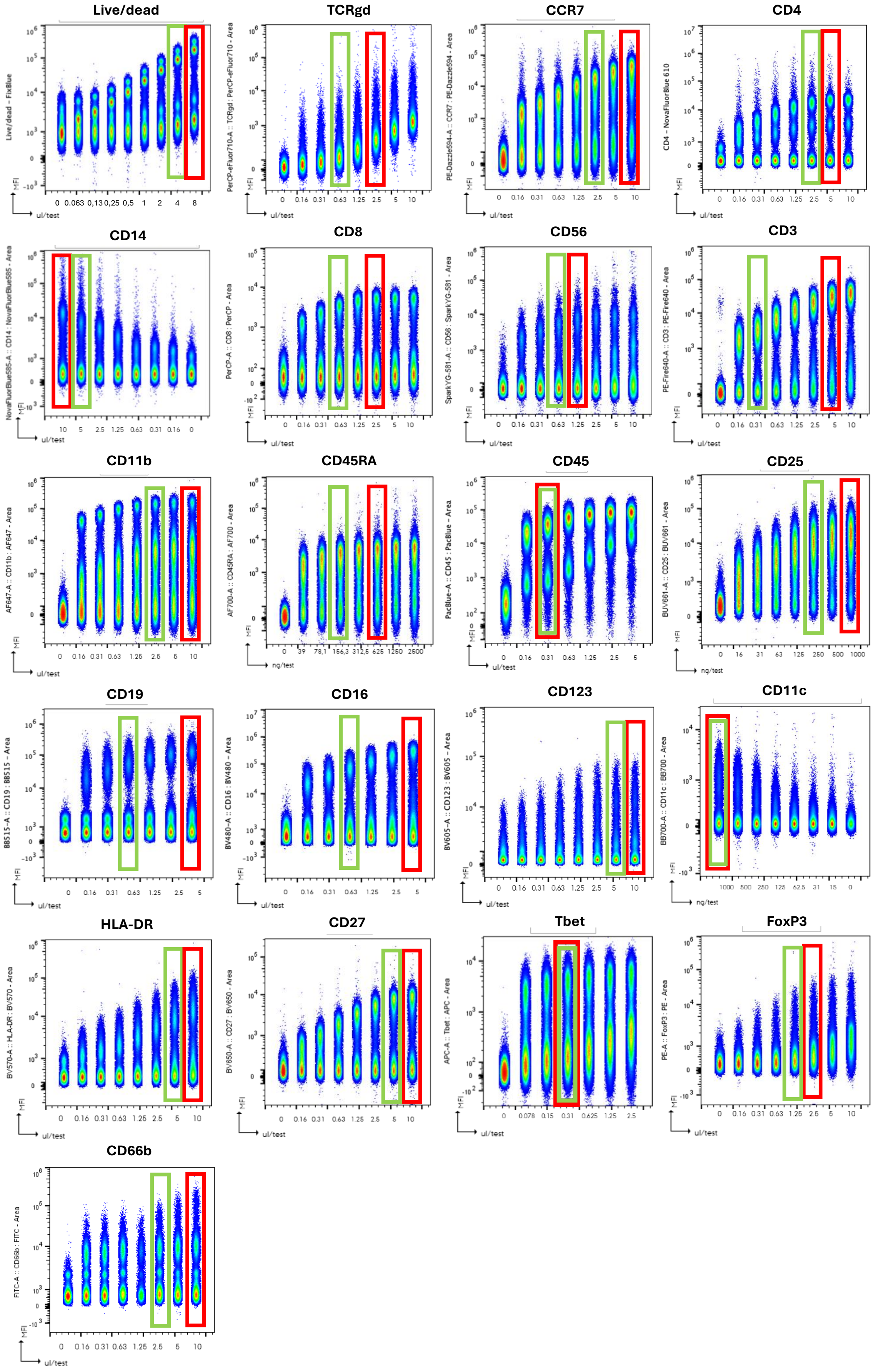

**Figure S3. Titration of all lineage and differentiation antibodies.** Example of antibody titration done on frozen PBMCs. The red square represents the recommended antibody concentration based on the calculated stain index, whereas the green square represents the selected antibody concentration used in the final panel.

Pedersen et al. figure S4

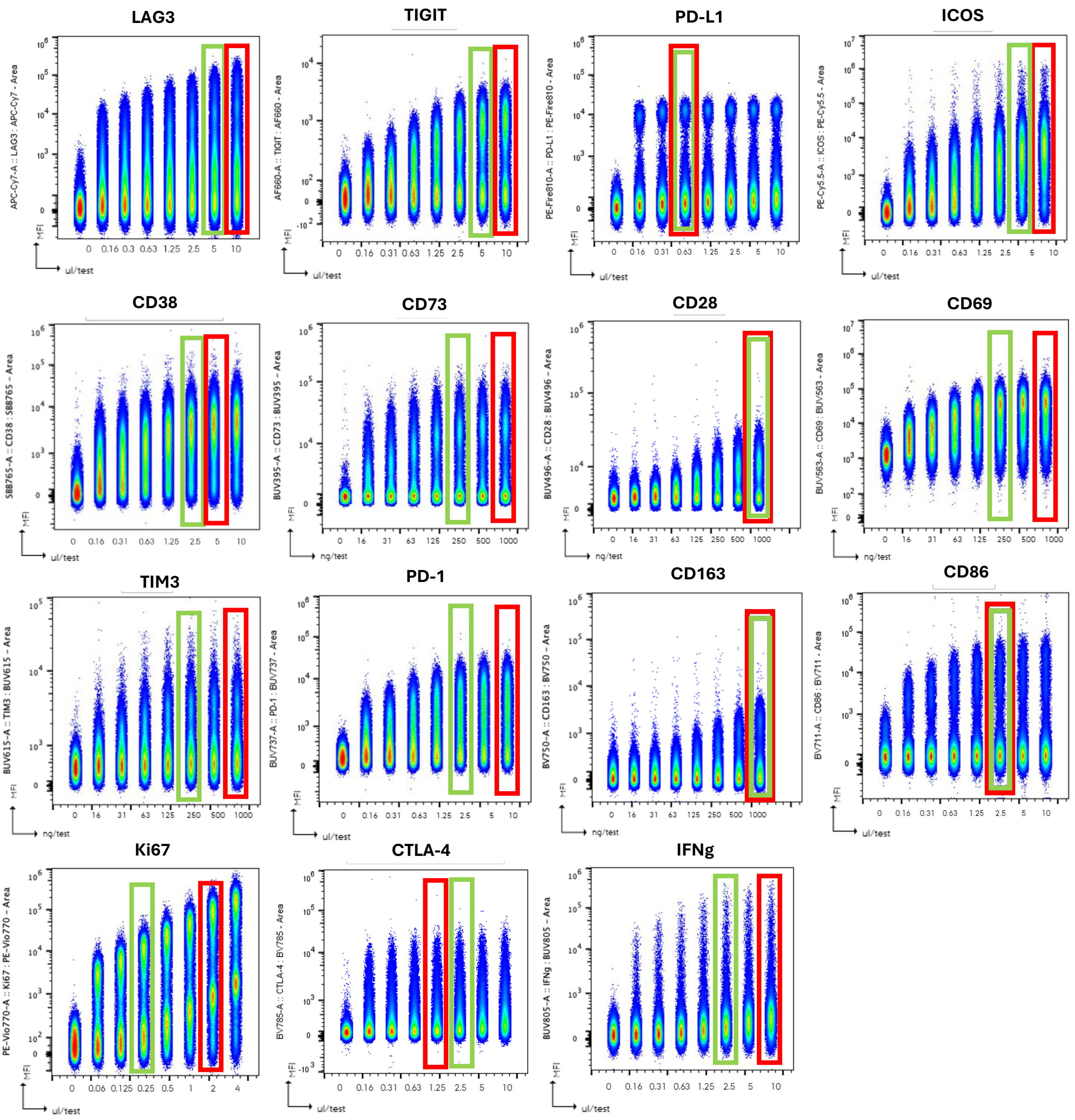

**Figure S4. Titration of all activation and exhaustion marker antibodies.** Example of antibody titration done on activated frozen PBMCs. The red square represents the recommended antibody concentration based on the calculated stain index, whereas the green square represents the selected antibody concentration used in the final panel

Pedersen et al. figure S5

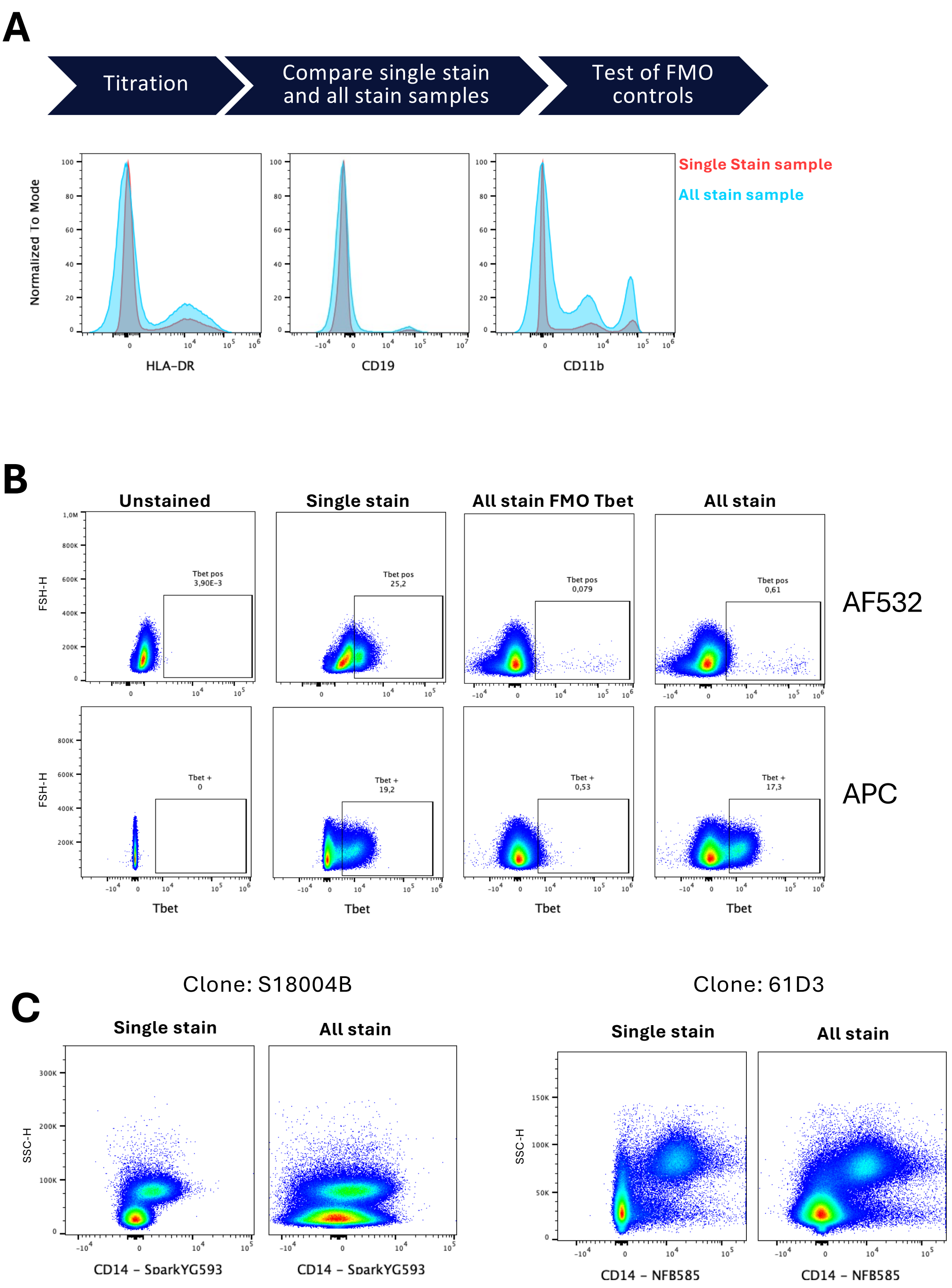

**Figure S5. Validation of antibody staining quality in all stained samples. A**) Schematic of the pipeline used for each antibody included in the final 37-marker panel. Examples of antibodies showing comparable staining quality between single stained and all stained samples. Single stain and all stain samples were performed on PBMCs, and histogram overlay plots were generated to compare staining quality between single stain and all stain samples for all markers. **B**) Example of poor antibody staining with Tbet AF532 which was improved by changing fluorochrome and antibody clone. Unstained, single stained, Fluoresence minus one (FMO) control, and all stained samples were prepared for Tbet AF532 (Clone 525803) and Tbet APC (Clone 4B10) on PBMCs to validate staining quality. **C**) Validation of CD14 antibodies in single stain and all stain samples. Poor separation between positive and negative population (single stain) and extensive data spreading (all stain) was seen for CD14 SparkYG593 (clone S18004B) (left hand side). Changing of the antibody to CD14 NFB585 (clone 61D3) significantly improved separation (single stain) and reduced data spread resulting in overall improved staining quality for the all stain sample.

Pedersen et al. figure S6

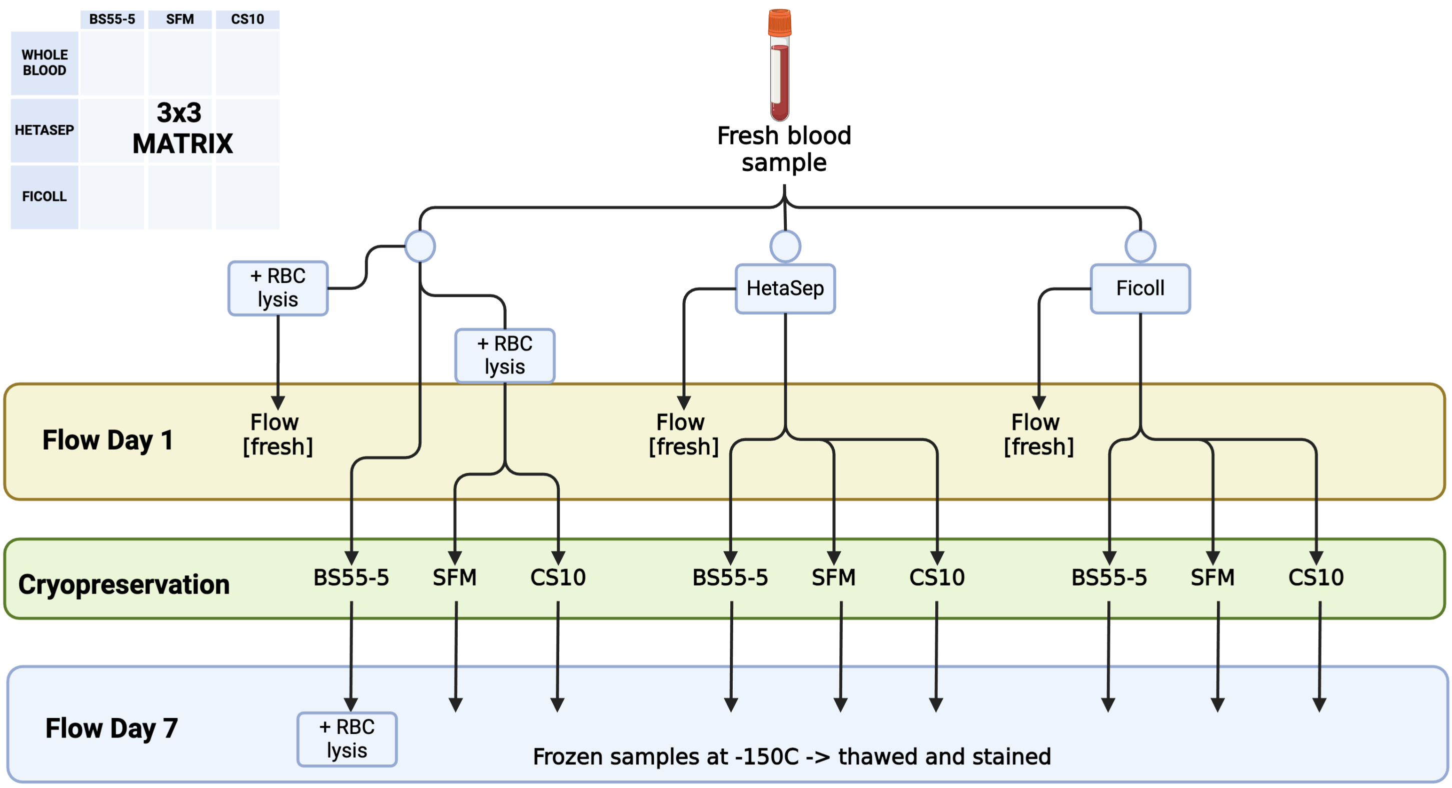

**Figure S6. The pipeline of sampling of liquid biopsies samples.**

Pedersen et al. figure S7

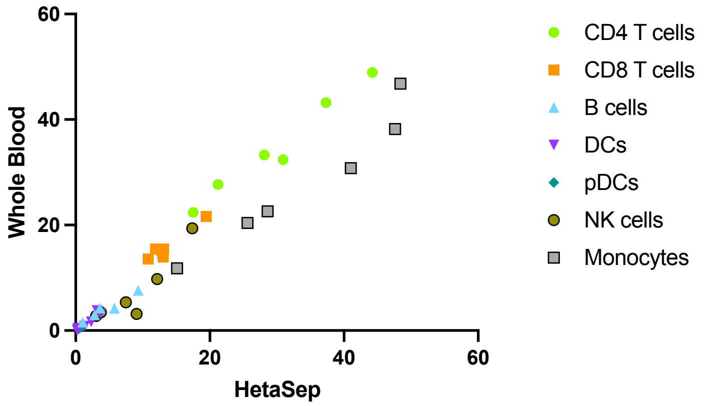

| Whole blood vs HetaSep | CD4 T cells | CD8 T cells | B cells | DCs | pDCs | NK cells | Monocytes |
|---|---|---|---|---|---|---|---|
| **Pearson r** | | | | | | | |
| r | 0,9845 | 0,9564 | 0,9579 | 0,9497 | 0,9877 | 0,9128 | 0,9682 |
| 95% conf. interval | 0,8603 to 0,9984 | 0,6473 to 0,9954 | 0,6576 to 0,9955 | 0,6023 to 0,9946 | 0,8875 to 0,9987 | 0,3908 to 0,9906 | 0,7308 to 0,9966 |
| R squared | 0,9692 | 0,9148 | 0,9176 | 0,9019 | 0,9755 | 0,8333 | 0,9373 |
| **P value** | | | | | | | |
| P (one-tailed) | 0,0002 | 0,0014 | 0,0013 | 0,0019 | 0,0001 | 0,0055 | 0,0008 |
| P value summary | *** | ** | ** | ** | *** | * | ** |

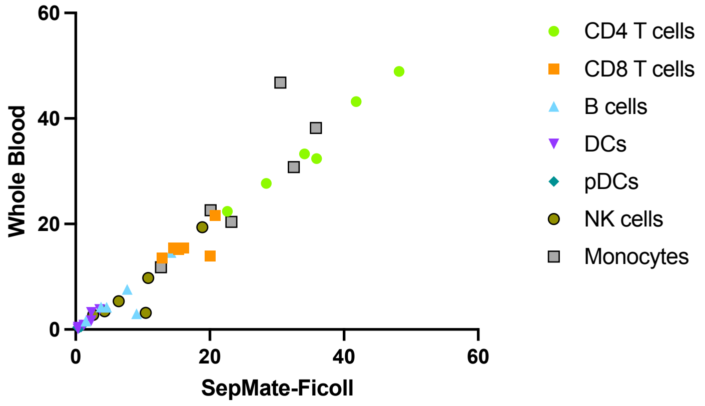

| Whole blood vs Ficoll | CD4 T cells | CD8 T cells | B cells | DCs | pDCs | NK cells | Monocytes |
|---|---|---|---|---|---|---|---|
| **Pearson r** | | | | | | | |
| r | 0,9865 | 0,6044 | 0,8495 | 0,9396 | 0,7042 | 0,8961 | 0,8511 |
| 95% confidence interval | 0,8774 to 0,9986 | -0,4066 to 0,9500 | 0,1222 to 0,9832 | 0,5391 to 0,9935 | -0,2506 to 0,9645 | 0,3099 to 0,9887 | 0,1279 to 0,9834 |
| R squared | 0,9732 | 0,3653 | 0,7217 | 0,8828 | 0,4959 | 0,8030 | 0,7244 |
| **P value** | | | | | | | |
| P (one-tailed) | 0,0001 | 0,1019 | 0,0161 | 0,0027 | 0,0592 | 0,0078 | 0,0158 |
| P value summary | *** | ns | * | ** | ns | ** | * |

**Figure S7. Correlation analysis on immune cell type frequencies between fresh whole blood and either HetaSep or Ficoll processed PBMCs.** The correlation plots included the immune cell type (CD4 T cells, CD8 T cells, B cells, DCs, pDCs, NK cells and Monocytes) frequencies as % of PBMCs.

### Pedersen et al. figure S8

| Whole blood vs BS55-5 | CD4 T cells | CD8 T cells | B cells | DCs | pDCs | NK cells | Monocytes |
|---|---|---|---|---|---|---|---|
| Pearson r | | | | | | | |
| r | 0,09696 | 0,3278 | 0,9636 | 0,5463 | 0,04054 | 0,5789 | 0,3986 |
| 95% conf. interval | -0,7756 to 0,8422 | -0,6591 to 0,9000 | 0,6976 to 0,9962 | -0,4766 to 0,9408 | -0,7973 to 0,8250 | -0,4389 to 0,9460 | -0,6105 to 0,9144 |
| R squared | 0,009401 | 0,1075 | 0,9285 | 0,2984 | 0,001644 | 0,3351 | 0,1588 |
| P value | | | | | | | |
| P (one-tailed) | 0,4275 | 0,2629 | 0,0010 | 0,1310 | 0,4696 | 0,1143 | 0,2169 |
| P value summary | ns | ns | *** | ns | ns | ns | ns |

| Whole blood vs CS10 | CD4 T cells | CD8 T cells | B cells | DCs | pDCs | NK cells | Monocytes |
|---|---|---|---|---|---|---|---|
| Pearson r | | | | | | | |
| r | 0,8848 | 0,7517 | 0,9430 | 0,9860 | -0,3435 | 0,7329 | 0,9611 |
| 95% conf. interval | 0,2599 to 0,9874 | -0,1536 to 0,9709 | 0,5600 to 0,9939 | 0,8727 to 0,9985 | -0,9033 to 0,6490 | -0,1941 to 0,9684 | 0,6800 to 0,9959 |
| R squared | 0,7829 | 0,5650 | 0,8892 | 0,9721 | 0,1180 | 0,5371 | 0,9238 |
| P value | | | | | | | |
| P (one-tailed) | 0,0096 | 0,0424 | 0,0024 | 0,0001 | 0,2525 | 0,0487 | 0,0011 |
| P value summary | ** | * | ** | *** | ns | * | ** |

| Whole blood vs SFM | CD4 T cells | CD8 T cells | B cells | DCs | pDCs | NK cells | Monocytes |
|---|---|---|---|---|---|---|---|
| Pearson r | | | | | | | |
| r | 0,5497 | 0,7776 | 0,9894 | 0,7057 | -0,2807 | 0,8743 | 0,7261 |
| 95% conf. interval | -0,4728 to 0,9413 | -0,09194 to 0,9743 | 0,9022 to 0,9989 | -0,2478 to 0,9647 | -0,8896 to 0,6875 | 0,2161 to 0,9861 | -0,2081 to 0,9675 |
| R squared | 0,3022 | 0,6047 | 0,9788 | 0,4980 | 0,07880 | 0,7644 | 0,5272 |
| P value | | | | | | | |
| P (one-tailed) | 0,1293 | 0,0343 | <0,0001 | 0,0586 | 0,2950 | 0,0114 | 0,0511 |
| P value summary | ns | * | **** | ns | ns | * | ns |

**Figure S8. Correlation analysis on immune cell type frequencies between fresh whole blood and each of the cryopreserved methods.** The correlation plots included the immune cell type (CD4 T cells, CD8 T cells, B cells, DCs, pDCs, NK cells and Monocytes) frequencies as % of PBMCs. Primary correlation plots can be seen in figure 4D.

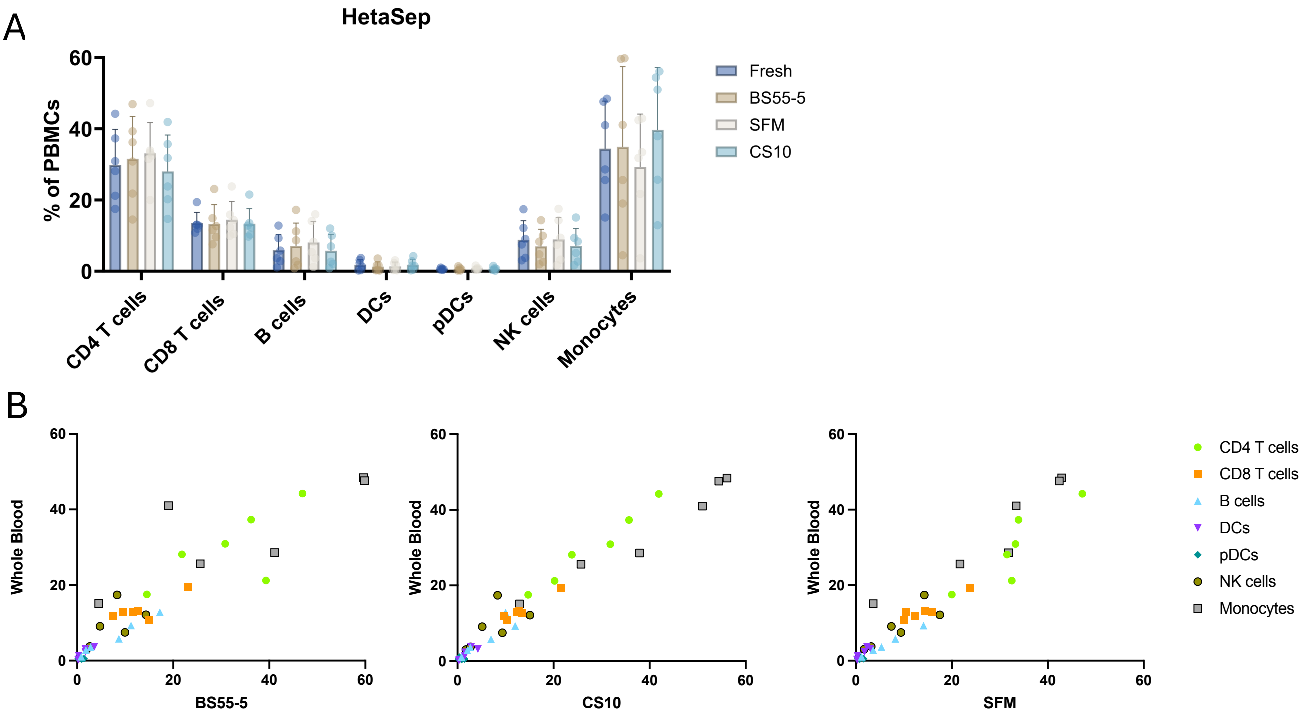

C

| Hetasep (fresh) vs Hetasep (BS55-5) | CD4 T cells | CD8 T cells | B cells | DCs | pDCs | NK cells | Monocytes |
|---|---|---|---|---|---|---|---|
| Pearson r | | | | | | | |
| r | 0,7042 | 0,8013 | 0,9854 | 0,9114 | -0,2468 | 0,6412 | 0,7976 |
| 95% conf. interval | -0,2506 to 0,9645 | -0,02935 to 0,9773 | 0,8675 to 0,9985 | 0,3836 to 0,9904 | -0,8818 to 0,7062 | -0,3551 to 0,9555 | -0,03955 to 0,9768 |
| R squared | 0,4959 | 0,6421 | 0,9709 | 0,8307 | 0,06093 | 0,4112 | 0,6362 |
| P value | | | | | | | |
| P (one-tailed) | 0,0592 | 0,0277 | 0,0002 | 0,0057 | 0,3186 | 0,0850 | 0,0286 |
| P value summary | ns | * | *** | ** | ns | ns | * |

| Hetasep (fresh) vs Hetasep (CS10) | CD4 T cells | CD8 T cells | B cells | DCs | pDCs | NK cells | Monocytes |
|---|---|---|---|---|---|---|---|
| Pearson r | | | | | | | |
| r | 0,9852 | 0,9784 | 0,9124 | 0,8880 | -0,2866 | 0,6527 | 0,9806 |
| 95% conf. interval | 0,8662 to 0,9984 | 0,8103 to 0,9977 | 0,3886 to 0,9905 | 0,2735 to 0,9877 | -0,8909 to 0,6841 | -0,3379 to 0,9572 | 0,8279 to 0,9980 |
| R squared | 0,9706 | 0,9573 | 0,8325 | 0,7885 | 0,08215 | 0,4260 | 0,9616 |
| P value | | | | | | | |
| P (one-tailed) | 0,0002 | 0,0003 | 0,0056 | 0,0091 | 0,2909 | 0,0800 | 0,0003 |
| P value summary | *** | *** | ** | ** | ns | ns | *** |

| Hetasep (fresh) vs Hetasep (SFM) | CD4 T cells | CD8 T cells | B cells | DCs | pDCs | NK cells | Monocytes |
|---|---|---|---|---|---|---|---|
| Pearson r | | | | | | | |
| r | 0,8810 | 0,9461 | 0,9874 | 0,9202 | -0,1454 | 0,8679 | 0,9458 |
| 95% conf. interval | 0,2437 to 0,9869 | 0,5796 to 0,9943 | 0,8852 to 0,9987 | 0,4292 to 0,9914 | -0,8560 to 0,7553 | 0,1907 to 0,9854 | 0,5777 to 0,9942 |
| R squared | 0,7762 | 0,8952 | 0,9750 | 0,8468 | 0,02114 | 0,7533 | 0,8946 |
| P value | | | | | | | |
| P (one-tailed) | 0,0102 | 0,0021 | 0,0001 | 0,0046 | 0,3917 | 0,0125 | 0,0022 |
| P value summary | * | ** | *** | ** | ns | * | ** |

**Figure S9. Changes in immune cell composition after cryopreservation of HetaSep isolated PBMCs.** A) Bar graph showing immune cell frequencies (as % of PBMCs) in fresh and cryopreserved HetaSep-isolated PBMCs. Statistical significance was tested with a RM one-way ANOVA test (n = 6). *$P < 0.05$. Bars indicate mean with SD as whiskers and individual donors as dots. B-C) Correlation plots with immune cell type frequencies as % of PBMCs in HetaSep-isolated PBMCs as compared to fresh whole blood.

Pedersen et al. figure S10

A

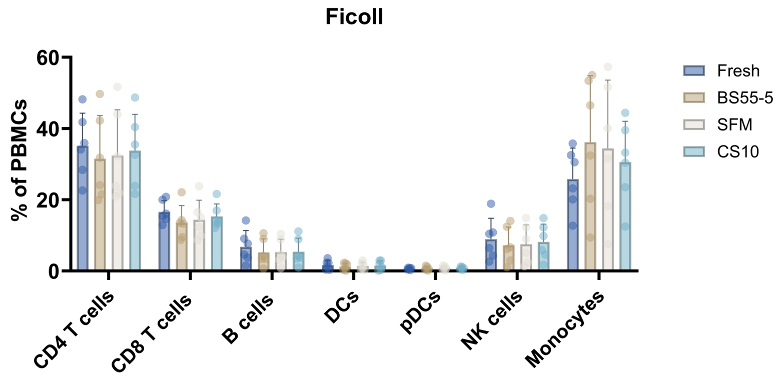

B

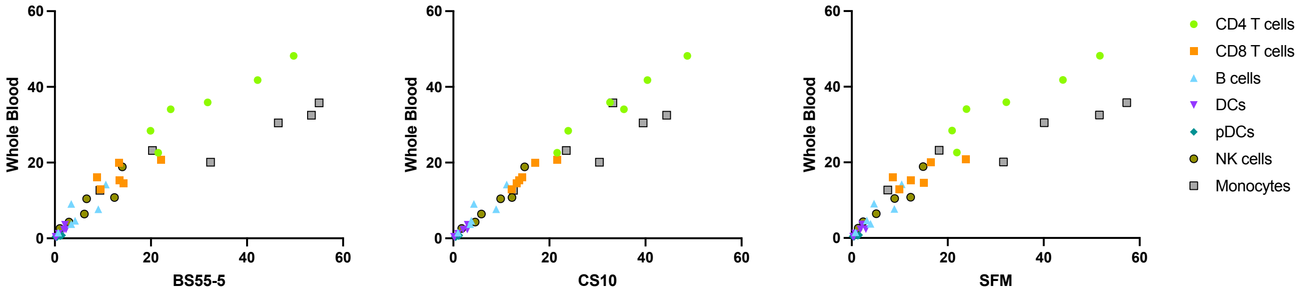

C

| Ficoll (fresh) vs Ficoll (BS55-5) | CD4 T cells | CD8 T cells | B cells | DCs | pDCs | NK cells | Monocytes |
|---|---|---|---|---|---|---|---|
| Pearson r | | | | | | | |
| r | 0,9375 | 0,6852 | 0,8243 | 0,9263 | 0,3514 | 0,9173 | 0,9442 |
| 95% conf. interval | 0,5265 to 0,9933 | -0,2847 to 0,9619 | 0,03858 to 0,9802 | 0,4623 to 0,9921 | -0,6437 to 0,9049 | 0,4139 to 0,9911 | 0,5673 to 0,9940 |
| R squared | 0,8789 | 0,4695 | 0,6795 | 0,8581 | 0,1235 | 0,8415 | 0,8914 |
| P value | | | | | | | |
| P (one-tailed) | 0,0029 | 0,0665 | 0,0218 | 0,0040 | 0,2473 | 0,0050 | 0,0023 |
| P value summary | ** | ns | * | ** | ns | ** | ** |

| Ficoll (fresh) vs Ficoll (CS10) | CD4 T cells | CD8 T cells | B cells | DCs | pDCs | NK cells | Monocytes |
|---|---|---|---|---|---|---|---|
| Pearson r | | | | | | | |
| r | 0,9788 | 0,9335 | 0,8775 | 0,9522 | 0,5291 | 0,9569 | 0,8386 |
| 95% conf. interval | 0,8132 to 0,9978 | 0,5031 to 0,9929 | 0,2290 to 0,9865 | 0,6191 to 0,9949 | -0,4950 to 0,9379 | 0,6506 to 0,9954 | 0,08454 to 0,9819 |
| R squared | 0,9580 | 0,8714 | 0,7700 | 0,9067 | 0,2800 | 0,9157 | 0,7032 |
| P value | | | | | | | |
| P (one-tailed) | 0,0003 | 0,0032 | 0,0108 | 0,0017 | 0,1402 | 0,0014 | 0,0185 |
| P value summary | *** | ** | * | ** | ns | ** | * |

| Ficoll (fresh) vs Ficoll (SFM) | CD4 T cells | CD8 T cells | B cells | DCs | pDCs | NK cells | Monocytes |
|---|---|---|---|---|---|---|---|
| Pearson r | | | | | | | |
| r | 0,9322 | 0,7881 | 0,8754 | 0,8438 | 0,4222 | 0,9548 | 0,9356 |
| 95% conf. interval | 0,4954 to 0,9927 | -0,06513 to 0,9756 | 0,2205 to 0,9863 | 0,1023 to 0,9825 | -0,5923 to 0,9189 | 0,6363 to 0,9952 | 0,5151 to 0,9931 |
| R squared | 0,8690 | 0,6211 | 0,7663 | 0,7120 | 0,1783 | 0,9117 | 0,8753 |
| P value | | | | | | | |
| P (one-tailed) | 0,0034 | 0,0313 | 0,0112 | 0,0173 | 0,2022 | 0,0015 | 0,0030 |
| P value summary | ** | * | * | * | ns | ** | ** |

**Figure S10. Changes in immune cell composition after cryopreservation of Ficoll isolated PBMCs.** A) Bar graph showing immune cell frequencies (as % of PBMCs) in fresh and cryopreserved Ficoll-isolated PBMCs. Statistical significance was tested with a RM one-way ANOVA test (n = 6). **P* < 0.05. Bars indicate mean with SD as whiskers and individual donors as dots. B-C) Correlation plots with immune cell type frequencies as % of PBMCs in HetaSep-isolated PBMCs as compared to fresh whole blood.

### Pedersen et al. figure S11

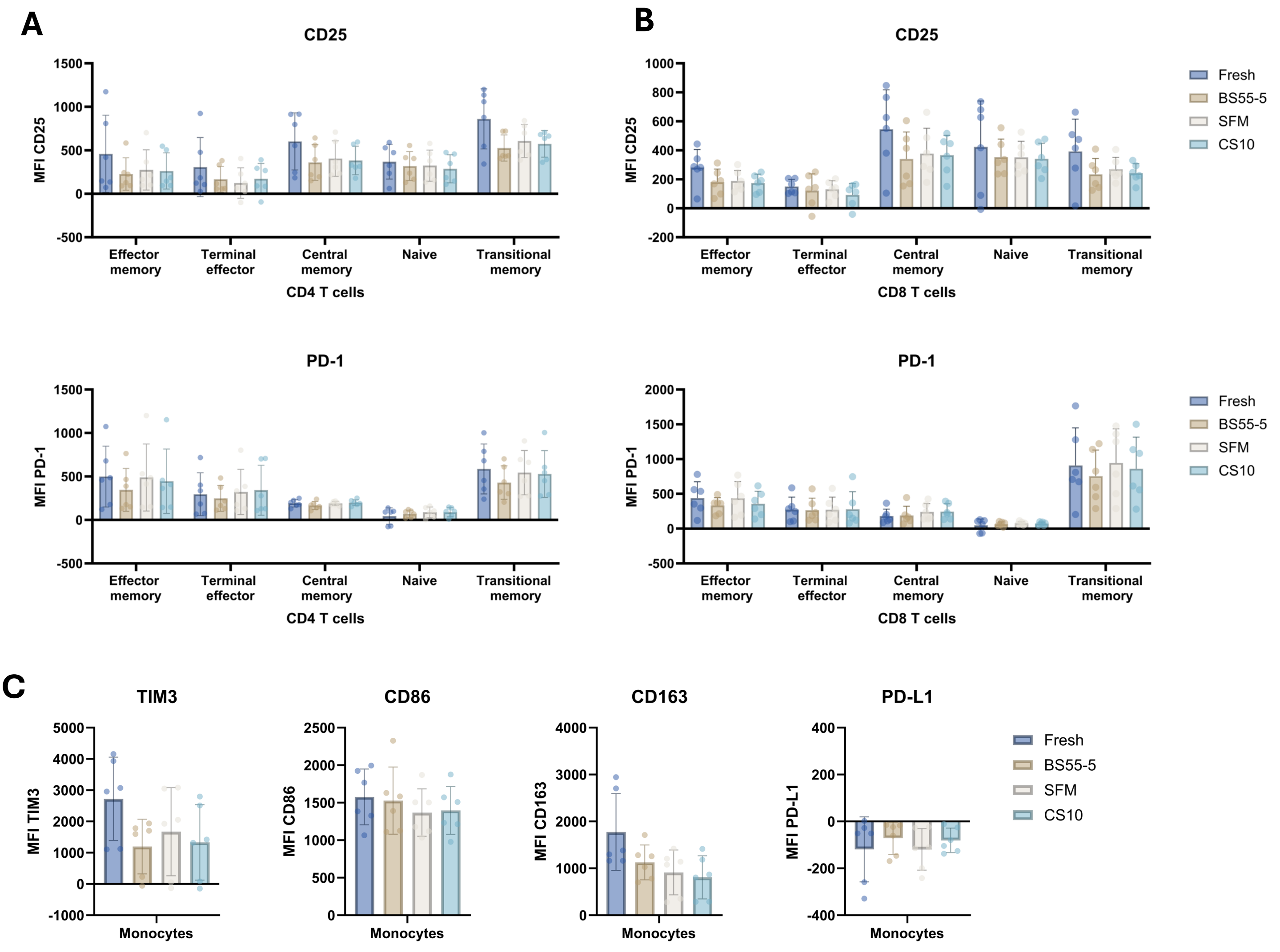

**Figure S11: Expression of functional markers in immune cells in whole blood samples.**
A) Sub-group analysis showing CD25 and PD-1 expression (MFI) on CD4 T cell subsets and B) CD8 T cell subsets across fresh and cryopreserved whole blood samples. C) Expression (MFI) of TIM3, CD86, CD163 and PD-L1 on Monocytes from fresh and cryopreserved samples. A-D) Statistical significance was tested using a RM one-way ANOVA with Bonferroni's multiple comparison test comparing each cryopreserved condition with fresh sample. *$P > 0.05$.

### Pedersen et al. figure S12

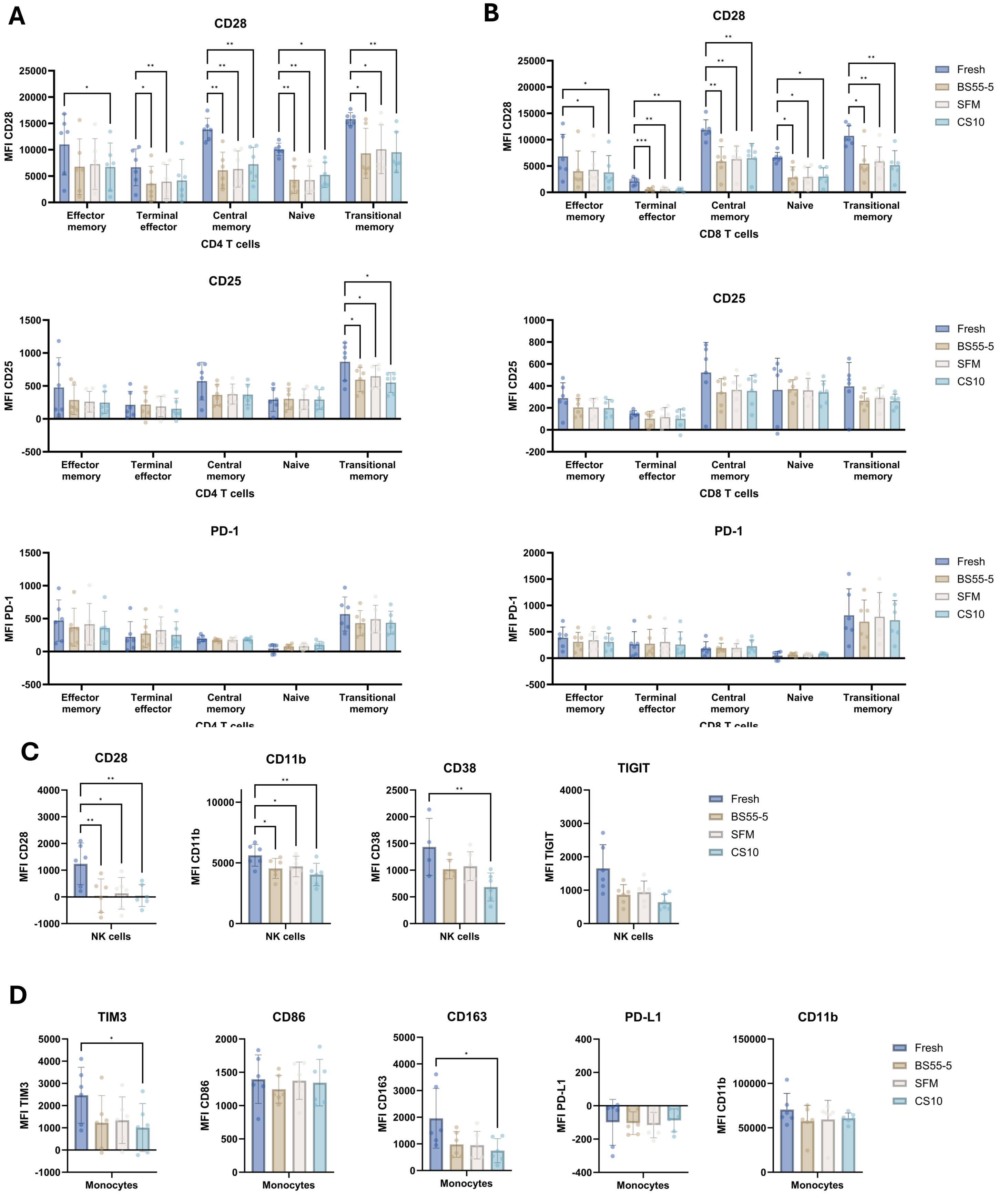

**Figure S12: Expression of functional markers in immune cells in HetaSep samples.**
A) Sub-group analysis showing CD28, CD25, and PD-1 expression (MFI) on CD4 T cell subsets and B) CD8 T cell subsets across fresh and cryopreserved HetaSep-isolated PBMCs. C) Expression (MFI) of CD28, CD11b, CD38, and TIGIT on NK cells from fresh and cryopreserved samples. D) Expression (MFI) of TIM3, CD86, CD163, PD-L1, and CD11b on Monocytes from fresh and cryopreserved samples. A-D) Statistical significance was tested using a RM one-way ANOVA with Bonferroni's multiple comparison test comparing each cryopreserved condition with fresh sample. **P* > 0.05 ; ***P* > 0.01.

### Pedersen et al. figure S13

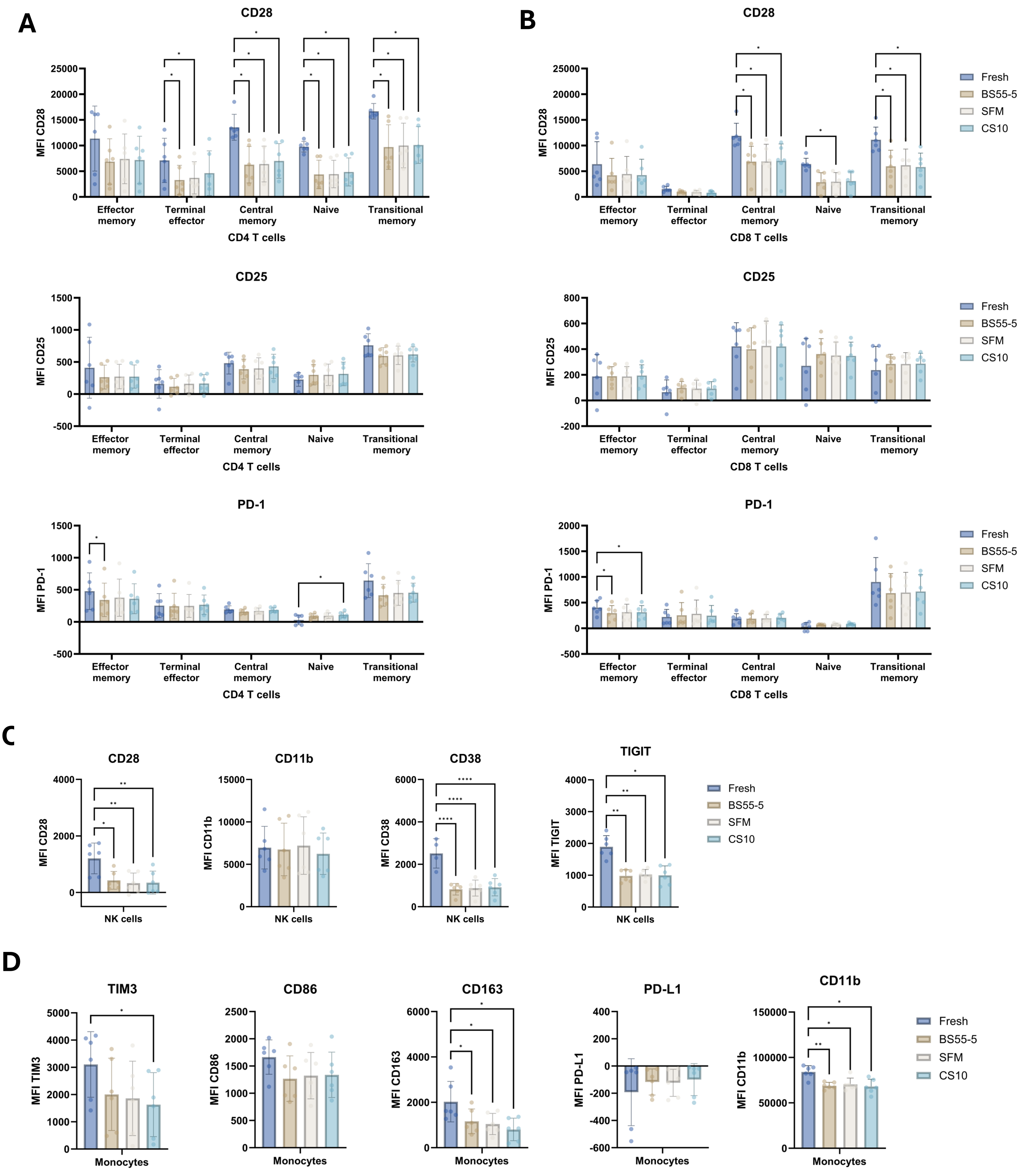

**Figure S13: Expression of functional markers in immune cells in Ficoll-gradient samples.**
A) Sub-group analysis showing CD28, CD25, and PD-1 expression (MFI) on CD4 T cell subsets and B) CD8 T cell subsets across fresh and cryopreserved Ficoll-isolated PBMCs. C) Expression (MFI) of CD28, CD11b, CD38, and TIGIT on NK cells from fresh and cryopreserved samples. D) Expression (MFI) of TIM3, CD86, CD163, PD-L1, and CD11b on Monocytes from fresh and cryopreserved samples. A-D) Statistical significance was tested using a RM one-way ANOVA with Bonferroni's multiple comparison test comparing each cryopreserved condition with fresh sample. *$P > 0.05$ ; **$P > 0.01$.

Pedersen et al. Table S1

| Live/dead mastermix | | |
|---|---|---|
| Reagent | Fluorochrome | Volume per sample (ul) |
| Live/dead | FixableBlue | 4 |
| PBS | | 96 |

| Mastermix 1 | | |
|---|---|---|
| Reagent | Fluorochrome | Volume per sample (ul) |
| TCRgd | PerCP-eFluor 710 | 0.625 |
| LAG3 | APC-Cy7 | 5 |
| TrueStain | | 5 |
| FACS wash | | 39.375 |

| Mastermix 2 | | |
|---|---|---|
| Reagent | Fluorochrome | Volume per sample (ul) |
| CCR7 | PE-Dazzle594 | 2.5 |
| CD4 | NovaFluorBlue 610 | 2 |
| CD14 | NovaFluorBlue 585 | 5 |
| CD66b | FITC | 2.5 |
| TIGIT | AF660 | 5 |
| PD-L1 | PE-Fire 810 | 0.63 |
| ICOS | PE-Cy5.5 | 5 |
| TrueStain | | 5 |
| CellBlox blocking buffer | | 5 |
| FACS wash | | 17.37 |

| Mastermix 3+4 | | |
|---|---|---|
| Reagent | Fluorochrome | Volume per sample (ul) |
| CD8 | PerCP | 0.63 |
| CD56 | Spark YG 581 | 1.25 |
| CD3 | PE-Fire 640 | 1.25 |
| CD11b | AF647 | 2.5 |
| CD45RA | AF700 | 1.25 |
| CD45 | Pacific Blue | 0.31 |
| CD38 | SBB765 | 2.5 |
| CD73 | BUV395 | 1.25 |
| CD28 | BUV496 | 5 |
| CD69 | BUV563 | 1.25 |
| TIM3 | BUV615 | 1.25 |
| CD25 | BUV661 | 1.25 |
| PD-1 | BUV737 | 2.5 |
| CD19 | BB515 | 0.63 |
| CD16 | BV480 | 0.63 |
| CD163 | BV750 | 5 |
| CD123 | BV605 | 5 |
| CD11c | BB700 | 5 |
| HLA-DR | BV570 | 5 |
| CD27 | BV650 | 5 |
| CD86 | BV711 | 2.5 |
| BD Brilliant stain buffer plus | | 10 |
| TrueStain monocyte blocker | | 5 |
| FACS wash | | 34.05 |

| Mastermix 5 | | |
|---|---|---|
| Reagent | Fluorochrome | Volume per sample (ul) |
| STING | BV421 | 2.5 |
| Ki67 | PE-Vio770 | 0.25 |
| Tbet | APC | 1.25 |
| FoxP3 | PE | 2.5 |
| CTLA4 | BV785 | 2.5 |
| IFNg | BUV805 | 2.5 |
| BD Brilliant stain buffer plus | | 10 |
| 1X Permeabilization buffer | | 28.5 |

**Table S1. Staining procedure for the 37-marker spectral flow panel.** The tables represent the volume of antibodies and other reagents used per sample in each of the five mastermixes in the 37-marker spectral flow panel.

### Pedersen et al. Table S2

| Specificity | Clone | Fluorophore | Company | Catalog number |
|---|---|---|---|---|
| CD73 | AD2 | BUV395 | BD bioscience | 742636 |
| CD28 | CD28.2 | BUV496 | BD bioscience | 741168 |
| CD69 | FN50 | BUV563 | BD bioscience | 748764 |
| TIM3 | 7D3 | BUV615 | BD bioscience | 752363 |
| CD25 | M-A251 | BUV661 | BD bioscience | 741608 |
| PD-1 | EH12.1 | BUV737 | BD bioscience | 612792 |
| STING | T3-680 | BV421 | BD bioscience | 564966 |
| CD16 | 3G8 | BV480 | BD bioscience | 566108 |
| CD163 | GHI/61 | BV750 | BD bioscience | 747185 |
| CD19 | HIB19 | BB515 | BD bioscience | 564456 |
| CD11c | 3.9 | BB700 | BD bioscience | 748270 |
| CD38 | AT13/5 | SBB765 | Bio-Rad | MCA1019SBB765 |
| CD45 | HI30 | Pacific Blue | Biolegend | 304029 |
| PD-L1 | 29E.2A3 | PE-Fire 810 | Biolegend | 329755 |
| LAG3 | 7H2C65 | APC-Cy7 | Biolegend | 369219 |
| CD123 | 6H6 | BV605 | Biolegend | 306026 |
| Ki67 | REA183 | PE-Vio770 | Miltenyi | 130-120-419 |
| HLA-DR | L243 | BV570 | Sony Biotechnology | 2138190 |
| CD27 | O323 | BV650 | Sony Biotechnology | 2114140 |
| CD86 | IT2.2 | BV711 | Sony Biotechnology | 2127200 |
| CTLA-4 | BNI3 | BV785 | Sony Biotechnology | 2448120 |
| CD66b | G10F5 | FITC | BioLegend | 305104 |
| CD8 | SK1 | PerCP | Sony Biotechnology | 2323540 |
| FoxP3 | 206D | PE | Sony Biotechnology | 2200540 |
| CD56 | QA17A16 | Spark YG 581 | Sony Biotechnology | 2562150 |
| CD3 | SK7 | PE-Fire 640 | Sony Biotechnology | 2324300 |
| CD11b | ICRF44 | AF647 | Sony Biotechnology | 2106595 |
| CCR7 | G043H7 | PE-Dazzle594 | Sony Biotechnology | 2366180 |
| CD45RA | HI100 | AF700 | Sony Biotechnology | 2120600 |
| IFNg | 4S.B3 | BUV805 | Thermo Fisher | 368-7319-42 |
| CD4 | SK3 | NFB610 | Thermo Fisher | H001T03B06 |
| TCRgd | B1.1 | PerCP-eFluor 710 | Thermo Fisher | 46-9959-42 |
| ICOS | ISA-3 | PE-Cy5.5 | Thermo Fisher | 35-9948-42 |
| TIGIT | MBSA43 | AF660 | Thermo Fisher | 606-9500-42 |
| CD14 | 61D3 | NovaFluor Blue 585 | Thermo Fisher | H019T03B04 |
| Tbet | 4B10 | APC | Thermo Fisher | 17-5825-82 |
| Viability | NA | Fixable blue | Thermo Fisher | L34962 |

**Table S2. List of antibodies included in the 37-marker spectral flow panel.** List of antibody name, clone, fluorochrome, manufacturer and catalogue number for each marker included in the 37-marker spectral flow panel.
